# Lysyl oxidase drives ccRCC progression by coordinating HIF-2α transcription program with tumor microenvironment

**DOI:** 10.64898/2026.07.24.739825

**Authors:** Burge Ulukan, Ozge Saatci, Ariel Madrigal, Minjun Kim, Wensheng Tian, Mustafa Soytas, Zohreh Mehrjoo, Ozlem Sener Sahin, Kukkamadi Sreenivas, Chintada Nageswara Rao, Tamiko Nishimura, Virginie Pillon, Jean-Sebastian Anoma, Elizabeth Hill, Janusz Rak, Campbell McInnes, Morag Park, Fadi Brimo, Simon Tanguay, Ryan Charles Russell, Hamed Najafabadi, Yasser Riazalhosseini, Ozgur Sahin

## Abstract

Clear cell renal cell carcinoma (ccRCC) is driven by persistent HIF-2α transcription program initiated by VHL loss, yet molecular mediators sustaining this program are poorly defined. Using single-cell transcriptomics, we identified lysyl oxidase (LOX) as a driver of ccRCC progression, selectively enriched in a hypoxia/epithelial-mesenchymal transition (EMT) gene program associated with poor outcome. While LOX oxidizes and stabilizes HIF-2α by antagonizing HUWE1-mediated ubiquitination and degradation, thereby sustaining HIF-2α-driven transcription in cancer cells, it also remodels extracellular matrix (ECM) and promotes angiogenesis in the tumor microenvironment (TME). Genetic or pharmacological inhibition of LOX destabilizes HIF-2α, disrupts ECM, inhibits angiogenesis, and suppresses tumor initiation, growth, and metastasis in vivo. LOX inhibition enhances anti-angiogenic therapy response and remains effective in belzutifan-resistant HIF-2α G323E-mutant tumors. Nuclear LOX protein correlates with nuclear HIF-2α in high-grade patient tumors. Together, LOX coordinates HIF-2α transcription program with TME and is a therapeutic target in ccRCC.

## Introduction

Clear cell renal cell carcinoma (ccRCC) is the most common subtype of kidney cancer, accounting for 75-85% of cases and the majority of kidney cancer-related mortality worldwide^1–3^. Although early-stage disease is often curable with surgical management or thermal ablation, advanced and metastatic ccRCC remains difficult to treat, with most patients eventually developing progressive disease resistant to standard-of-care therapies^4,5^. These clinical challenges underscore a critical need to define the molecular mechanisms that sustain tumor progression and therapeutic resistance in ccRCC and identify novel targets to effectively treat this highly aggressive disease.

At the genomic level, ccRCC is driven by early and recurrent loss of chromosome 3p, which harbors multiple tumor suppressor genes, including *VHL, PBRM1, SETD2,* and *BAP1.* Among these, VHL is inactivated in approximately 80% of ccRCC tumors, with additional recurrent mutations observed in *PBRM1* (∼40%), *SETD2* (∼20%), and *BAP1* (∼15%)^6,7^. Notably, inactivation of VHL, an E3 ubiquitin ligase, is the canonical initiating event, resulting in constitutive stabilization of hypoxia-inducible factor 2-alpha (HIF-2α) under normoxic conditions^8^. While VHL loss initiates HIF-2α stabilization, how HIF-2α proteostasis is sustained and activates its transcriptional program within these highly heterogenous VHL-deficient tumors during tumor progression remains poorly understood. Indeed, large-scale single-cell and spatial transcriptomic studies, including our recent work, reveal that this pseudohypoxic state is not uniform^9^. Rather, it reflects a continuum of malignant cell states characterized by varying degrees of pseudohypoxia-associated transcriptional activity within a hypervascular and stiff extracellular matrix (ECM)^10–13^. This spatial and cellular heterogeneity suggests that sustained HIF-2α signaling may be differentially regulated across different tumor cells and regions, contributing to tumor progression. However, little is known about the tumor-intrinsic mechanisms that selectively sustain HIF-2α activity within aggressive malignant cell states in ccRCC.

Upregulation of vascular endothelial growth factor (VEGF)^14^ is among the major downstream targets of HIF-2α, promoting angiogenesis and tumor progression. Along these lines, VEGF/VEGFR-targeted tyrosine kinase inhibitors (TKIs) have become a cornerstone of ccRCC therapy and further improved clinical outcomes in combination with immune checkpoint inhibitors^15^. However, the clinical benefit of these therapies is limited due to systemic toxicities and the rapid emergence of adaptive resistance^16,17^. Persistent HIF-2α activity in ccRCC tumors has led to the development of HIF-2α inhibitors^18–20^, including belzutifan^21^ and next-generation HIF-2α antagonists, such as NKT2152^20^, and casdatifan^22^. While the FDA-approved belzutifan, a direct HIF-2α dimerization blocker, demonstrated clinical activity^23^, the duration of response is relatively limited, and is associated with on-target toxicities, including severe anemia, hypoxia, and fatigue that frequently necessitate treatment interruption or dose modification^24^. Moreover, resistance-conferring mutations within the HIF-2α PAS-B pocket (e.g., G323E) impair drug binding^25,26^, while co-occurring genomic alterations (e.g., *TP53* mutations) further compromise therapeutic responses^27,28^. Considering the hypervascular and highly heterogeneous nature of ccRCC, there is an urgent need to identify novel therapeutic vulnerabilities to simultaneously target both the tumor and its microenvironment in order to improve clinical outcomes. Upregulation of vascular endothelial growth factor (VEGF)^14^ is among the major downstream targets of HIF-2α, promoting angiogenesis and tumor progression. Along these lines, VEGF/VEGFR-targeted tyrosine kinase inhibitors (TKIs) have become a cornerstone of ccRCC therapy and further improved clinical outcomes in combination with immune checkpoint inhibitors^15^. However, the clinical benefit of these therapies is limited due to systemic toxicities and the rapid emergence of adaptive resistance^16,17^. Persistent HIF-2α activity in ccRCC tumors has led to the development of HIF-2α inhibitors^18–20^, including belzutifan^21^ and next-generation HIF-2α antagonists, such as NKT2152^20^, and casdatifan^22^. While the FDA-approved belzutifan, a direct HIF-2α dimerization blocker, demonstrated clinical activity^23^, the duration of response is relatively limited, and is associated with on-target toxicities, including severe anemia, hypoxia, and fatigue that frequently necessitate treatment interruption or dose modification^24^. Moreover, resistance-conferring mutations within the HIF-2α PAS-B pocket (e.g., G323E) impair drug binding^25,26^, while co-occurring genomic alterations (e.g., *TP53* mutations) further compromise therapeutic responses^27,28^. Considering the hypervascular and highly heterogeneous nature of ccRCC, there is an urgent need to identify novel therapeutic vulnerabilities to simultaneously target both the tumor and its microenvironment in order to improve clinical outcomes.

Here, we identify lysyl oxidase (LOX), a copper-dependent amine oxidase best known for collagen and elastin crosslinking^29,30^ in the tumor microenvironment, as a driver of ccRCC progression. Mechanistically, we show that HIF-2α is a novel non-canonical substrate of LOX, which stabilizes HIF-2α by antagonizing its ubiquitin-dependent proteasomal degradation, thereby sustaining downstream hypoxia-associated transcriptional programs. Therapeutically, LOX inhibition simultaneously disrupts both HIF-2α-driven gene programs and tumor microenvironment remodeling, leading to tumor growth inhibition, metastasis suppression, and overcoming resistance to standard-of-care therapies used in ccRCC.

## Results

### Single-cell RNA profiling identifies a LOX-enriched hypoxia-EMT-associated malignant cell program, associated with disease progression in ccRCC tumors

To systematically identify malignant cell-intrinsic transcriptional programs associated with ccRCC progression, we analyzed 59 intra-tumoral gene expression programs reported in Madrigal et al., derived from malignant cells using single-cell transcriptomes across 19 renal cell carcinoma (RCC) specimens (**Fig. 1A**). Among these, Gene Program 60 (GP60) emerged as one of the most clinically relevant malignant programs^31^. GP60 showed higher activity in ccRCC compared to other major subtypes and was associated with worse clinical outcome among malignant gene programs in the TCGA-KIRC cohort **(Fig. 1B)**. Consistent with the pseudohypoxic biology of VHL-deficient ccRCC, GP60 showed strong enrichment for hypoxia-related gene sets and a significant overlap with HIF-2α transcriptional signatures (**Fig. 1A, C**).

**Figure 1:**
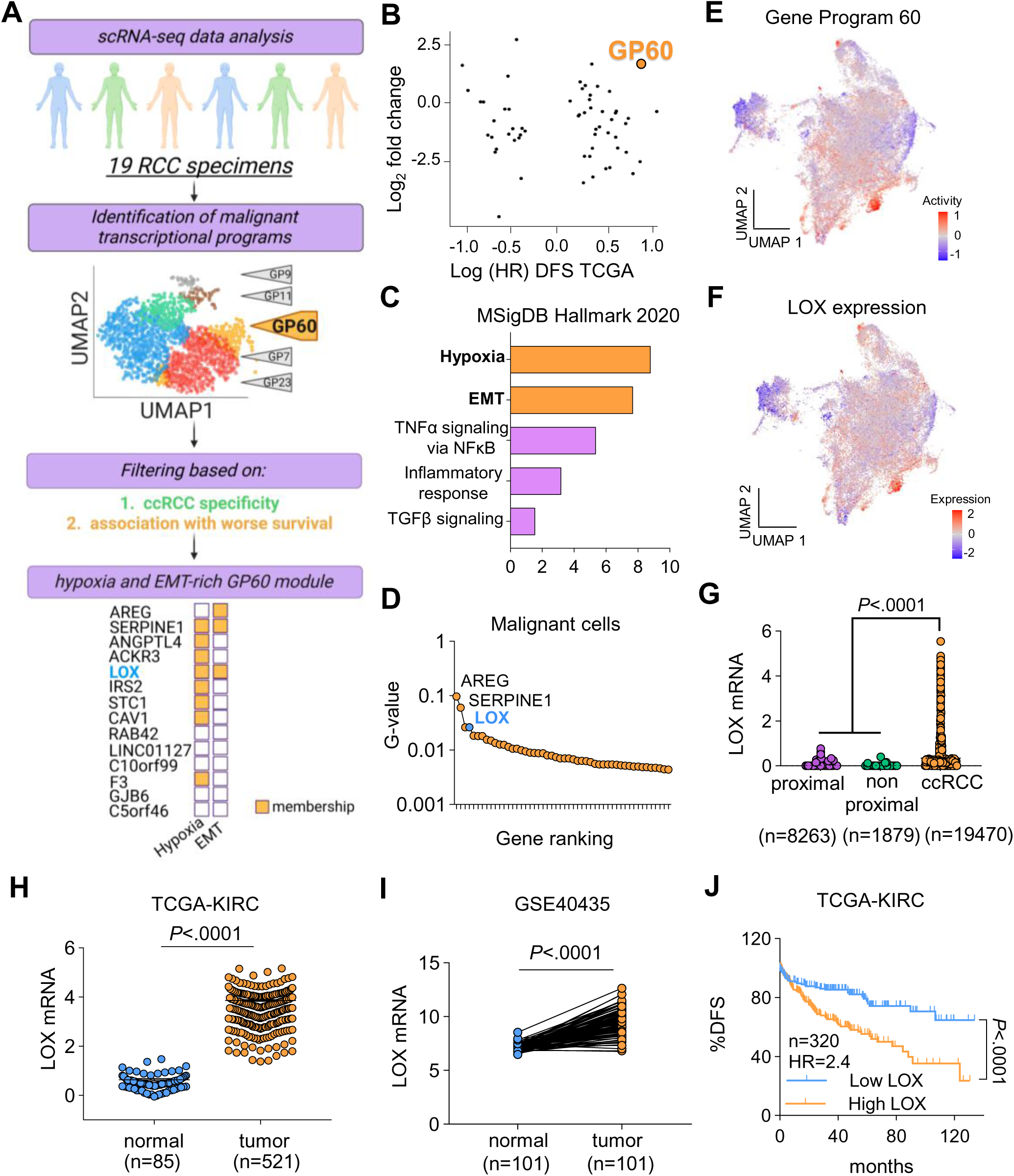
Single-cell RNA profiling of ccRCC tumors identifies a LOX-enriched hypoxia-EMT-associated malignant cell program, associated with disease progression. **(A)** Schematic workflow leading to the identification of GP60 in malignant renal cell carcinoma cells^31^. Following filtering based on ccRCC specificity and association with poor clinical outcome, GP60 emerged as a hypoxia- and EMT-associated malignant gene program. The gene list shown represents membership of GP60 top genes in hypoxia (including the Hallmark hypoxia gene (MsigDB^62^) and HIF-derived gene signatures^31^), and EMT. Created with BioRender.com. **(B)** Scatterplot ranking gene expression programs by ccRCC subtype specificity and association with disease-free survival (DFS). The x-axis shows log hazard ratio for DFS in the TCGA-KIRC cohort; the y-axis shows mean log_2_ standardized fold change comparing GP60 activity in ccRCC vs. other subtypes, computed using the scoreMarkers function from scran^63^.^63^. **(C)** Pathway enrichment analysis of GP60 genes using MSigDB Hallmark pathways, revealing significant enrichment for hypoxia and epithelial-mesenchymal transition (EMT) signatures. **(D)** Gene ranking of *LOX* among the top members of GP60. The x-axis shows the G value, a non-negative weight representing the relative contribution of each gene to program 60 (weights for all genes in GP60 sum to 1). **(E)** UMAP embedding shows GP60 activity in malignant RCC cells. The embedding was generated from the activities of 59 intra-tumoral gene expression programs identified in malignant RCC cells^31^. **(F)** UMAP embedding shows the normalized latent-log expression of *LOX* in malignant RCC cells. **(G)** Single-cell expression analysis showing enrichment of *LOX* in ccRCC (n = 19470) compared to proximal (n = 8263) and non-proximal normal (n = 1879) kidney epithelial cells (GSE152938). **(H)** *LOX* expression in ccRCC tumors (n = 521) and normal kidney tissues (n = 85) from the TCGA-KIRC cohort. **(I)** *LOX* expression in matched ccRCC tumor vs. adjacent normal kidney tissues from the GSE40435 cohort (n = 101). **(J)** Kaplan-Meier curve showing DFS in TCGA-KIRC patients (n = 320 HR 2.4), stratified by high vs. low *LOX* expression (median cutoff). Statistical significance for pairwise comparisons was assessed using paired two-sided Student’s *t*-test (I) or Mann-Whitney *U* test (H), as appropriate. Survival differences were evaluated using the log-rank (Mantel-Cox) test (J). *P*-values are indicated.

Within GP60, Serpin Family E member 1 (*SERPINE1*) and *LOX* emerged as the top-ranked genes associated with hypoxia and EMT-associated signatures (**Fig. 1A, D**). Given the established role of SERPINE1 in ccRCC progression^32^, we focused on LOX. Notably, we observed an almost one-to-one overlap of LOX and GP60 activity across malignant cells (**Fig. 1E, F**). Analysis of an independent single cell RNA-seq (scRNA-seq) dataset^33,34^, confirmed that LOX expression is enriched in malignant cells compared to normal proximal tubular epithelial and non-proximal cells (**Fig. 1G).** Consistent with this, bulk RNA-seq analysis showed upregulation of LOX in RCC tumors of the TCGA kidney renal clear cell carcinoma (KIRC) cohort (**Fig. 1H**) and of an independent ccRCC dataset (GSE40435) compared to normal kidney tissue (**Fig. 1I**). Importantly, higher LOX expression was associated with significantly worse DFS in TCGA-KIRC (**Fig. 1J**), consistent with the prognostic impact of GP60^31^, supporting a clinically relevant role for LOX in ccRCC progression.

### LOX stabilizes HIF-2α through lysine oxidation and antagonizing its HUWE1-mediated ubiquitination

We next sought to define how LOX functionally relates to hypoxia-driven transcriptional states in ccRCC. Our scRNA-seq data analysis showed that LOX expression strongly correlates with GP60 (**Fig. 2A**) and with HIF-2α (**Fig. 2B**) activities, but not HIF-1α transcriptional activity (**Supplementary Fig. 1A**), within malignant cells. We therefore asked whether LOX contributes to the maintenance of HIF-2α-driven programs beyond serving solely as a downstream target of HIF-2α in ccRCC^35,36^. To address this, we leveraged intronic and exonic *LOX* expression as proxies for nascent transcription and mature, protein-producing transcripts, respectively. Notably, exonic *LOX* expression, reflecting mature protein-producing transcripts, showed a stronger correlation with EMT, hypoxia, and HIF-2α transcriptional signatures than intronic reads across single cells (**Fig. 2C, D**), indicating that LOX protein (i.e., output of exonic reads), rather than *LOX* mRNA alone, is associated with sustained HIF-2α activity, supporting an upstream regulatory role of LOX on HIF-2α.

**Figure 2:**
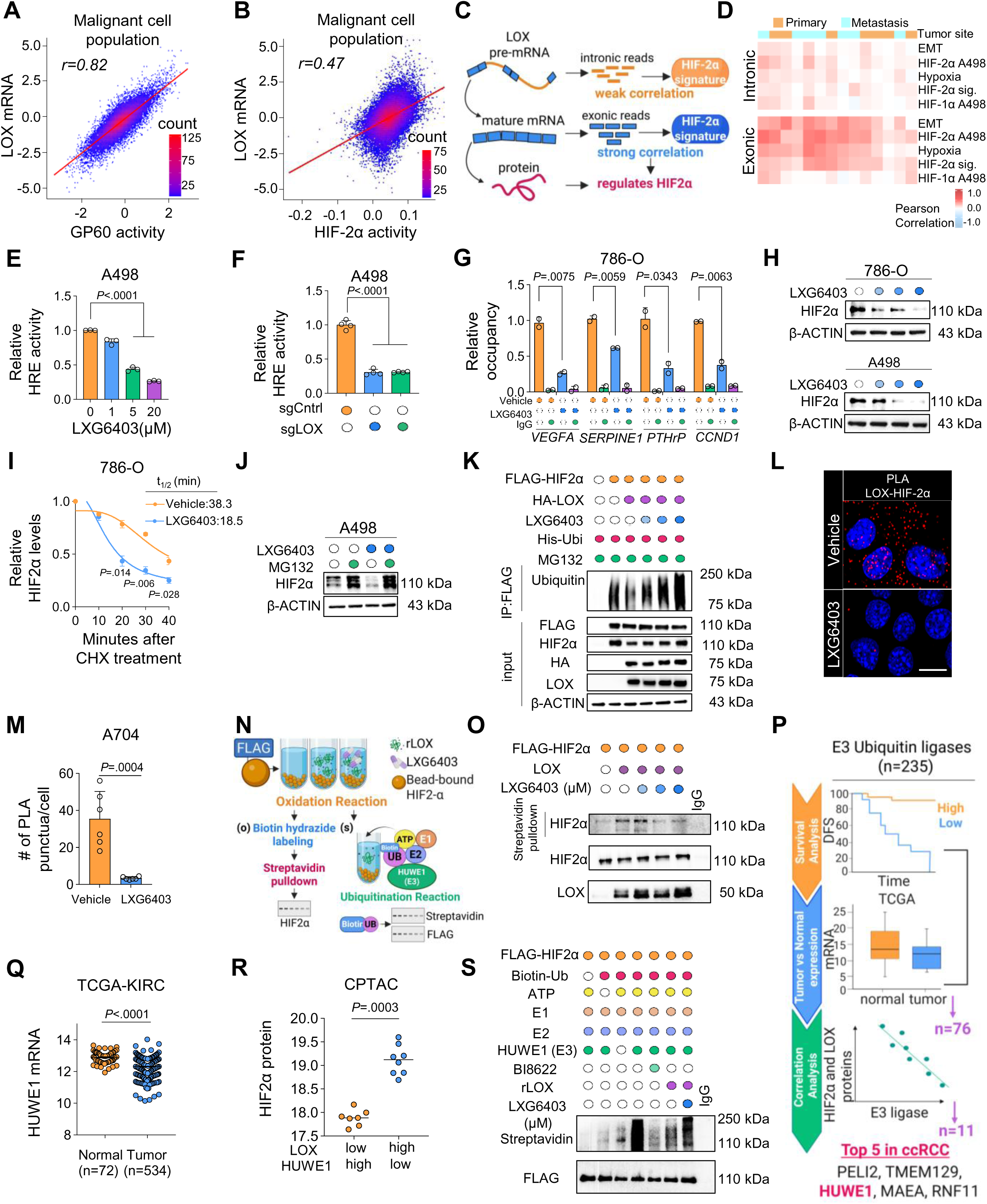
LOX stabilizes HIF-2α through lysine oxidation and antagonizing HUWE1-mediated ubiquitination. (A, **B)** Relationship between *LOX* expression, GP60 activity and HIF-2α activity in malignant RCC cells. Left: scatterplot showing latent-log expression of *LOX* (x-axis) vs. GP60 activity (y-axis). The line represents a linear model fit, and Pearson correlation values are indicated. Right: same as left but comparing *LOX* expression to estimated HIF-2α activity (See Methods). **(C)** Schematic illustrating the inferred relationship between intronic and exonic mRNA levels of *LOX* and HIF-2α activity. Higher correlation between intronic read counts and HIF2α activity would be consistent with HIF-2α acting upstream of *LOX* transcription, whereas higher correlation between exonic *LOX* read counts and HIF-2α activity would support an association between mature LOX transcripts and HIF-2α activity. Created with BioRender.com. **(D)** Relationship between normalized exonic and intronic LOX expression and activity of selected gene sets in ccRCC malignant cells. Heatmap showing Pearson correlation values for each sample (columns). Top bar indicates metastasis status. Gene sets were derived from the Hallmark MsigDB collection, a HIF-2α signature and HIF-derived gene signatures^31^. **(E, F)** HRE-GFP reporter assay indicating reduced HIF-2α transcriptional activity in 786-O cells in response to dose-dependent inhibition of LOX (n = 3 technical replicates) (E) or CRISPR-mediated LOX knockout (F) (n = 4 technical replicates). **(G)** HIF-2α ChIP-qPCR analysis showing reduced HIF-2α binding to *VEGFA*, *SERPINE1*, *PTHrP,* and *CCND1* promoters upon LOX inhibition (n = 2). **(H)** Immunoblot analysis shows dose-dependent reduction of HIF-2α protein levels in 786-O and A498 cells following LOX inhibition with LXG6403. β-ACTIN was used as a loading control here and in all immunoblots unless otherwise indicated. **(I)** Quantification of HIF-2α half-life from CHX chase experiments from untreated (control) and LXG6403 (30 μM for 16 hours) treated samples (n = 2). **(J)** Immunoblot analysis of HIF-2α levels in 786-O cells treated with vehicle or LXG6403 (30 μM) in the presence or absence of the proteasome inhibitor MG132, indicating proteasome-dependent regulation. **(K)** Ubiquitination assay showing reduced HIF-2α ubiquitination upon LOX overexpression (HA-LOX) and increased ubiquitination following LOX inhibition with LXG6403 for 12 hours, MG132 (10 μM for 6 hours) (n = 3). Increasing blue color intensity corresponds to increasing LXG6403 concentrations (7.5, 15, and 30 μM). **(L)** Representative proximity ligation assay (PLA) in A704 cells demonstrating nuclear proximity between LOX and HIF-2α, and reduction upon LXG6403 treatment (30 μM for 6 hours) Scale bar = 25 μm. (n = 2). **(M)** Quantification of PLA puncta per nucleus from (l) (n = 6 different fields). **(N)** Experimental workflow of in vitro oxidation assay followed by in vitro ubiquitination. Created with BioRender.com. **(O)** In vitro oxidation assay using bead-bound HIF-2α as substrate and recombinant LOX protein, followed by biotin-hydrazide labeling and streptavidin pulldown, demonstrating direct LOX-mediated oxidation on HIF-2α and reduced lysine oxidation upon LOX inhibition (LXG6403 5 μM, 15 μM and 30 μM) (n = 3). **(P)** The analysis pipeline to identify potential downstream effector E3 ligases that are involved in LOX inhibition-mediated HIF-2α degradation. Created with BioRender.com. **(Q)** Levels of *HUWE1* mRNA in ccRCC tumor (n = 534) vs. normal (n = 72) from TCGA. **(R)** Levels of HIF-2α in ccRCC tumors with low LOX/high HUWE1 vs. high LOX/low HUWE1. **(S)** In vitro ubiquitination assay using bead-bound HIF-2α as substrate and HUWE1-enriched lysates as the E3 ligase source, demonstrating that LOX-mediated oxidation suppresses HUWE1-dependent HIF-2α ubiquitination, while LOX inhibition (LXG6403 15 μM) restores ubiquitination in a cell-free system (n = 3). Data are shown as mean ± SD for biological experiments, unless otherwise indicated. Patient-derived data are shown as mean ± SEM. Statistical significance was determined using unpaired two-tailed Student’s *t*-test (G, M, Q, R) or one-way ANOVA with Dunnett’s multiple-comparison corrections (E, F). *P*-values are indicated. Colors denote experimental groups and are used consistently across graphs and representative images throughout the manuscript unless otherwise indicated. Open circles indicate control conditions, whereas filled circles indicate experimental conditions.

To test the hypothesis that LOX is an upstream regulator of HIF-2α transcriptional program, we first performed a HIF-responsive element (HRE)-GFP reporter assay which showed marked reduction in HIF-2α transcriptional activity upon LOX inhibition (**Fig. 2E, F Supplementary Fig. 1B**). Concordantly, qRT-PCR confirmed downregulation of canonical HIF-2α target mRNAs, including *VEGFA*, *CCND1*, as well as *LOX* and *LOXL2*, following CRISPR-mediated LOX knockout (**Supplementary Fig. 1C**), consistent with the inhibition of HIF-2α-dependent transcriptional programs. To determine the direct effects of LOX on HIF-2α-mediated transcription of its target genes, we performed ChIP-qPCR. LOX inhibition significantly reduced the HIF-2α occupancy on the promoter regions of *VEGFA, SERPINE1, PTHrP,* and *CCND1* (**Fig. 2G**).

At the protein level, LOX inhibition led to a dose-dependent reduction in HIF-2α abundance (**Fig. 2H**), without altering *EPAS1* mRNA (gene name for HIF-2α) levels (**Supplementary Fig. 1D**), nominating a post-translational regulation as the potential mode of action. Notably, LOX inhibition closely phenocopied the effects of VHL add-back on lowering HIF-2α protein levels and reducing clonogenicity in A498 cells (**Supplementary Fig. 1E-G**). To test the effects of LOX on HIF-2α protein stability, we performed cycloheximide (CHX) chase experiment which revealed reduced HIF-2α protein half-life upon LOX inhibition (**Fig. 2I, Supplementary Fig. 2A**). Proteasome inhibition with MG132 restored HIF-2α levels, whereas lysosomal inhibition with bafilomycin A1 had no effect (**Fig. 2J, Supplementary Fig. 2B**), implicating ubiquitin-proteasome-mediated degradation. Consistent with this mechanism, HIF-2α ubiquitination increased upon LOX inhibition in a dose-dependent manner and was suppressed upon LOX overexpression (**Fig. 2K**). Proximity ligation assay (PLA) showed a robust nuclear and cytoplasmic interaction between LOX and HIF-2α that was significantly diminished upon LOX inhibition (**Fig. 2L, M)**. Reciprocal co-immunoprecipitation confirmed a physical interaction between LOX and HIF-2α that was reduced following LXG6403 treatment (**Supplementary Fig. 2C, D**). Notably, chromatin-associated HIF-2α was also reduced upon LOX inhibition (**Supplementary Fig. 2E**), consistent with impaired HRE activity (**Fig. 2E, F Supplementary Fig. 1B)**.

Given that LOX is an amine oxidase^37^, we next asked if HIF-2α could be a direct enzymatic substrate of LOX. In vitro oxidation assays using bead-bound FLAG-HIF-2α incubated with recombinant LOX showed robust lysine oxidation detected by biotin-hydrazide labeling that was abolished by LOX inhibition in a dose-dependent manner (**Fig. 2N, left, O**). This result was further validated in a fully cell-free system using recombinant GST-HIF-2α, confirming direct LOX-dependent oxidation, independent of bead-bound or cellular components, as shown by the OxyBlot analysis, and similarly suppressed by LXG6403 (**Supplementary Fig. 2F**).

Since both lysine oxidation and ubiquitination occur on the ε-amino group lysine residues, we hypothesized that LOX-mediated oxidation may interfere with ubiquitin conjugation at modified lysine sites on HIF-2α. To this end, we first sought to identify the E3 ubiquitin ligase responsible for HIF-2α turnover in VHL-deficient ccRCCs. We performed an integrative analysis of major E3 ligase families by combining (i) associations with patient survival, (ii) differential expression between ccRCC tumors and normal kidney, and (iii) inverse correlation with HIF-2α and LOX protein expression. This analysis identified 11 candidate E3 ligases, with the top-ranked candidates including PELI2, TMEM129, HUWE1, MAEA, and RNF11 (**Fig. 2P**). Among those, HUWE1 emerged as the most mechanistically relevant E3 ligase; unlike RING-type E3 ligases that function primarily as scaffolds, HUWE1 is a HECT-domain E3 ligase that directly transfers ubiquitin to substrate lysine residues^38^, providing a biochemical rationale for LOX-mediated interference. Consistent with this, high HUWE1 expression was significantly associated with improved patient survival (**Supplementary Fig. 2G**), and HUWE1 expression was reduced in ccRCC tumors compared to normal kidney tissues (**Fig. 2Q**). In addition, tumors characterized by low LOX and high HUWE1 protein expression exhibited markedly lower HIF-2α protein levels compared to tumors with high LOX and low HUWE1 expressions (**Fig. 2R**), supporting a potential antagonistic relationship between LOX and HUWE1. Consistent with a role for HECT-domain ligases in this pathway, we showed that pharmacological inhibition of HECT ligases using heclin^39^ restored HIF-2α stability upon LOX inhibition (**Supplementary Fig. 2H**).

To determine whether LOX-mediated HIF-2α oxidation directly interferes with the HUWE1-dependent ubiquitination, pre-oxidized or control bead-bound HIF-2α was subjected to in vitro ubiquitination assays using HUWE1-immunoprecipitated lysate as the E3 source (**Fig. 2N, right)**. HUWE1 inhibition markedly reduced HIF-2α ubiquitination in a cell-free system, demonstrating its functionality. Notably, LOX-mediated HIF-2α oxidation interferes with HUWE1-dependent ubiquitination, whereas LOX inhibition restored HIF-2α ubiquitination (**Fig. 2S**). Together, these findings suggest that LOX directly oxidizes lysine residues on HIF-2α, thereby antagonizing HUWE1-mediated ubiquitination and proteasomal degradation, contributing to sustained HIF-2α transcriptional activity in VHL-deficient ccRCCs.

### LOX is crucial for ccRCC tumor growth

Since LOX-mediated lysine oxidation canonically drives collagen crosslinking and integrin signaling^40,41^, we first assessed the effects of LOX inhibition on cell viability in three-dimensional collagen-matrigel culture. Both genetic (sgLOX) and pharmacological (LXG6403) inhibition of LOX led to a decrease in 3D cell viability in a dose-dependent manner (in the case of LXG6403) (**Fig. 3A-F**). Notably, these effects were more pronounced in *VHL*-mutant lines compared to *VHL*-wild type ACHN and CAKI-1 cells (**Supplementary Fig. 3A, B**).

**Figure 3:**
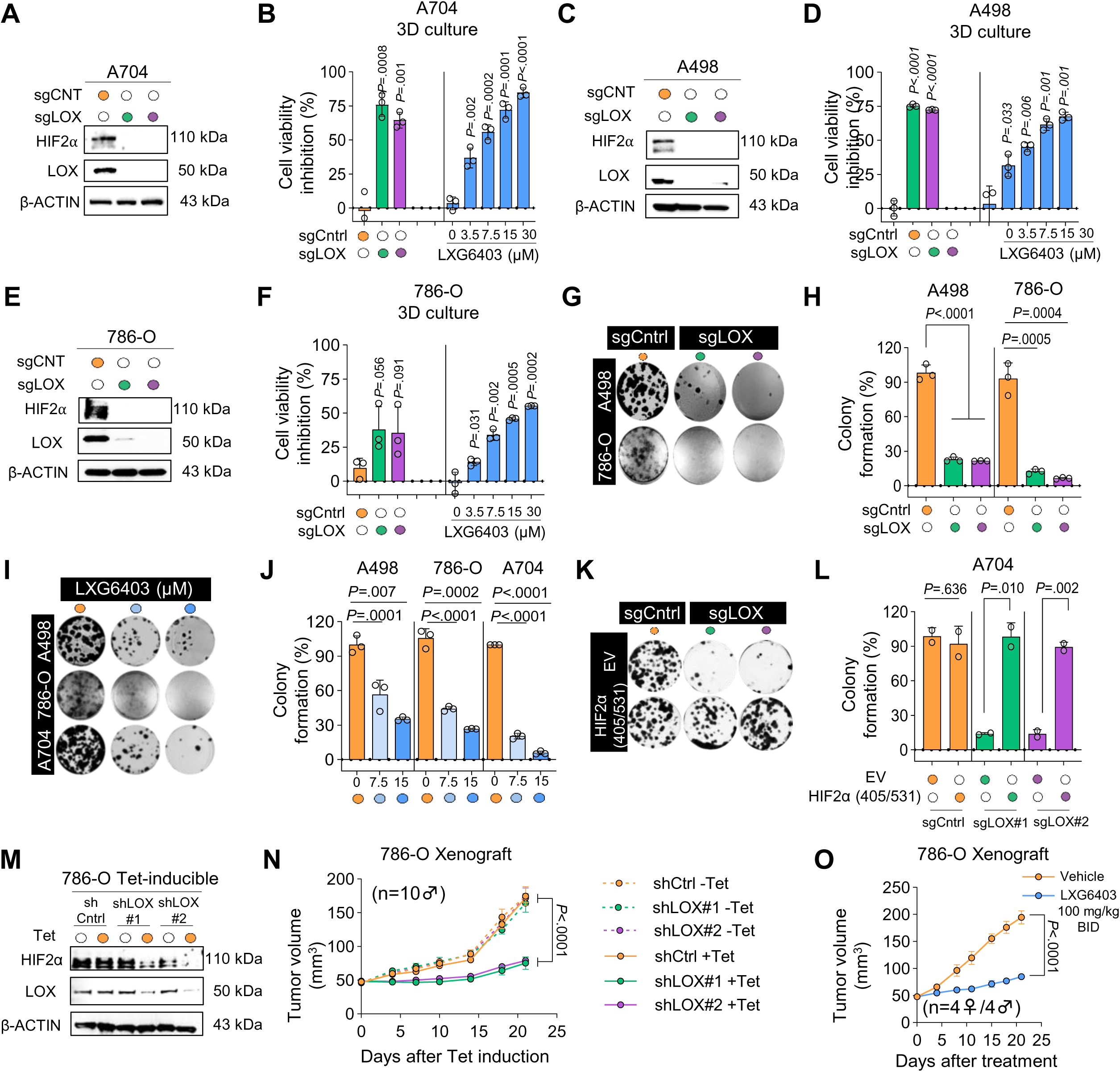
LOX is crucial for ccRCC tumor growth. **(A)** Immunoblot validation of CRISPR-mediated LOX knockout using two independent sgRNAs in A704. **(B)** Collagen-matrigel-embedded 3D viability assays showing growth inhibition upon LOX knockout or pharmacological inhibition in A704 cells (n = 3). **(C)** Immunoblot validation of CRISPR-mediated LOX knockout using two independent sgRNAs in A498. **(D)** Collagen-matrigel-embedded 3D viability assays showing growth inhibition upon LOX knockout or pharmacological inhibition in A498 cells (n = 2). **(E)** Immunoblot validation of CRISPR-mediated LOX knockout using two independent sgRNAs in 786-O. **(F)** Collagen-matrigel-embedded 3D viability assays showing growth inhibition upon LOX knockout or pharmacological inhibition in 786-O cells (n = 2). **(G, H)** Colony formation assay showing reduced clonogenic potential in A498 (G) and 786-O (H) cells upon LOX knockout with two independent sgRNAs (n = 3). **(I, J)** Pharmacological LOX inhibition with LXG6403 (7.5μM and 15 µM) decreases colony formation in A498, 786-O, and A704 cells (I) and its quantification (J) (n = 3). (**K, L**) Colony formation assay showing a reduced clonogenic potential in sgLOX cells and rescue upon overexpression of degradation-resistant HIF-2α (P405/P531A) (K) and its quantification (L) (n = 2). (**M)** Immunoblot of sh-inducible 786-O cells confirming the knockdown. **(N)** Growth curves of subcutaneous 786-O xenografts expressing inducible shLOX (tet-on system) or non-targeting control (shCtrl) in NSG male mice (n = 10). Tetracycline induction (1 mg/ml in drinking water) suppresses tumor growth relative to shCtrl. **(O)** Sex-balanced 786-O xenograft (n = 8; 4 female/4 male NSG mice) cohort treated with vehicle or LXG6403 (100 mg/kg, BID, p.o.), showing >50% tumor growth suppression relative to vehicle. Data are shown as mean ± SD for biological experiments, unless otherwise indicated. In vivo data are shown as mean ± SEM. An unpaired two-sided Student’s t-test (L) was used to calculate the statistical difference between the two groups. One-way ANOVA with Dunnett’s test (B, D, F, H, J) or Holm’s (N, O) was performed to compare more than one group. *P*-values are indicated.

To test the effects of cell-centric LOX on clonogenic growth, we performed loss-of-function studies in *VHL*-mutant ccRCC cell lines with high LOX expression. CRISPR-mediated knockout using two different sgRNAs targeting LOX (sgLOX) effectively depleted LOX and significantly reduced the clonogenic potential of 786-O and A498 cells (**Fig. 3C, E, G, H**). Consistent with this, pharmacological inhibition with the small-molecule inhibitor of LOX, LXG6403^42^, mimicked genetic inhibition and induced a dose-dependent decrease in clonogenic potential of cells (**Fig. 3I, J**). To determine whether this effect is mediated through HIF-2α stability, we re-expressed a degradation-resistant *HIF2A* mutant (P405A/P531A)^43^ in sgLOX cells (**Supplementary Fig. 3C**). Restoration of degradation-resistant HIF-2α rescued colony formation to near control levels (**Fig. 3K, L**), supporting HIF-2α as a key effector of LOX.

We next tested if LOX is required for ccRCC tumor growth in vivo. We first utilized a tetracycline-inducible LOX knockdown system (shLOX) in 786-O cells, which reduced LOX and HIF-2α protein levels (**Fig. 3M**). These cells were then injected into mice, and LOX knockdown was induced by tetracycline administration once tumors became palpable (∼50 mm^3^). LOX inhibition significantly suppressed tumor growth compared to non-targeting controls (**Fig. 3N**) with no significant changes in body weight (**Supplementary Fig. 4A**), showing that LOX is required for the growth of ccRCC tumors. Next, we tested the therapeutic potential of targeting LOX using LXG6403 in vivo and demonstrated a significant suppression of tumor growth in a sex-balanced cohort of the same model (**Fig. 3O**), phenocopying shLOX-mediated tumor suppression, with no significant body weight changes **(Supplementary Fig. 4B)**. Together, these data demonstrate that VHL-deficient ccRCC cells depend on LOX for survival, clonogenic potential, and tumor growth, and highlight LOX as a therapeutically targetable vulnerability in ccRCC.

### LOX promotes tumor cell plasticity, self-renewal, and metastasis in ccRCC

Given the strong enrichment of the EMT signature in the GP60 hypoxic malignant cell state, where LOX is among the top-ranked genes overlapping with program activity (**Fig. 1**), we tested the effects of LOX on EMT in ccRCC. Genetic depletion of LOX (sgLOX) or pharmacological inhibition with LXG6403 resulted in a significant reduction in the expression of mesenchymal markers (CDH2, Vimentin, Slug, and ZEB-1) and an upregulation of epithelial markers (CDH1 and ZO-1) at both mRNA (**Fig. 4A, Supplementary Fig. 5A**) and protein levels (**Fig. 4B, Supplementary Fig. 5B**). This was accompanied by impaired cell migration and invasion, in both sgLOX-expressing (**Fig. 4C, D**) and LXG6403-treated cells (**Fig. 4E, F, Supplementary Fig. 5C, D**).

**Figure 4:**
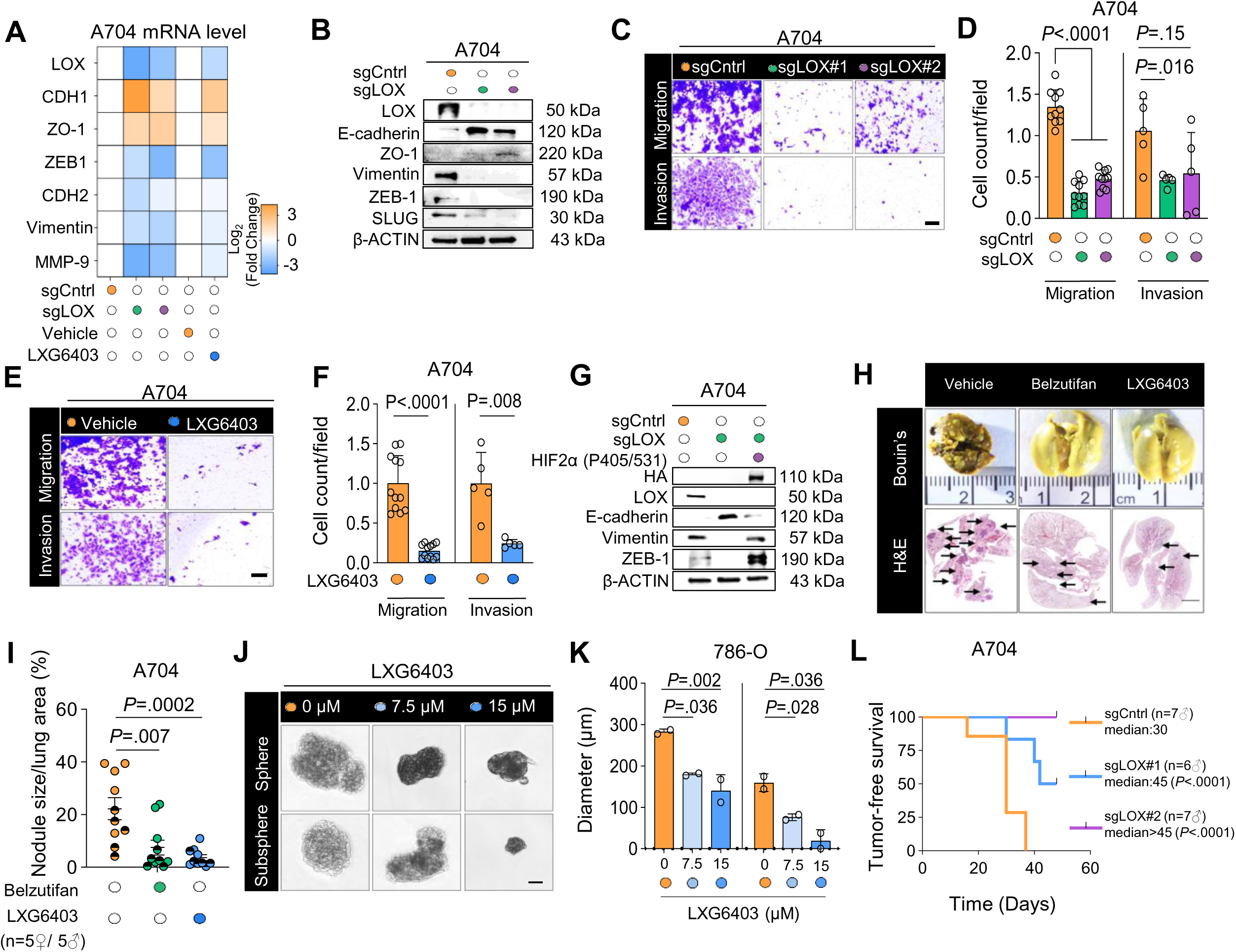
LOX promotes tumor cell plasticity, self-renewal, and metastasis programs in ccRCC. **(A)** qRT-PCR analysis showing downregulation of mesenchymal markers and upregulation of epithelial markers upon LOX inhibition (n = 2). **(B)** Immunoblot confirming EMT marker changes in sgLOX-expressing A704 cells (n = 2). **(C-F)** Representative images and quantification of migration (24 h) (n = 10-12 different images) and invasion (48 h) (n = 5 different images) in sgLOX-expressing (C, D) and LXG6403-treated (30 µM) A704 cells (E, F) (n = 2). Scale bar = 100 μm. **(G)** Immunoblot showing restoration of mesenchymal markers (Vimentin and ZEB1) and suppression of the epithelial marker E-cadherin upon overexpression of degradation-resistant HIF-2α in sgLOX cells. **(H)** Representative Bouin’s-fixed and H&E-stained lung sections from mice injected subcutaneously with A704 cells, showing metastatic nodules (arrows and circles) across treatment groups (vehicle, LXG6403 (100 mg/kg, BID, p.o.), or HIF-2α inhibitor (Belzutifan, 3mg/kg, BID, p.o.) (n = 10 (5F/5M) NSG mice). Scale bar = 500 μm. **(I)** Quantification of lung metastatic burden from (H), expressed as the number of metastatic nodules per lung area (n = 10 (5F/5M)). Graphs display color-coded treatment groups, with female mice shown as solid symbols and male mice as half-colored symbols. **(J, K)** Representative images of spheroids (J) and quantification of diameter of spheres/sub-spheres (K) in 786-O cells treated with LXG6403 (7.5μM and 15 µM) (n = 2). Scale bar = 200 µm. Data are shown mean ± SD. **(L)** Tumor-free survival in n = 6-7 NSG male mice implanted with A704 cells expressing sgCtrl or sgLOX. Tumor initiation is defined as the first measurement ≥40 mm³. Data are shown as mean ± SD for biological experiments, unless otherwise indicated. In vivo data are shown as mean ± SEM. Curves were compared by the log-rank (Mantel-Cox) test (L). An unpaired two-sided Student’s t-test (A, F) or One-way ANOVA with Dunnett’s test (D, I, K) were used to calculate the statistical difference between the two groups. *P*-values are indicated.

To determine whether LOX-driven EMT programs depend on HIF-2α, we re-expressed degradation-resistant HIF-2α in LOX-deficient cells. Restoration of HIF-2α in sgLOX cells rescued EMT markers (**Fig. 4G**), indicating that HIF-2α mediates LOX-dependent regulation of tumor cell plasticity. To determine the effects of LOX inhibition-mediated suppression of HIF-2α and downstream effects on in vivo metastasis, we injected highly metastatic A704 cells subcutaneously into NSG mice and treated the mice with LXG6403 or the HIF-2α inhibitor, belzutifan. Both agents significantly reduced metastatic burden, as evidenced by fewer lung nodules per section on Bouin’s and H&E staining (**Fig. 4H, I**), without significant changes in body weight (**Supplementary Fig. 5E**).

Given that EMT has been shown to generate stem-like tumor-initiating cells^44^, we next asked if LOX-dependent EMT programs also regulate self-renewal and tumor initiation in ccRCC. LOX inhibition significantly impaired tumor sphere formation ability of 786-O cells by reducing both sphere diameter and number in a dose-dependent manner (**Fig. 4J, K, Supplementary Fig. 5F**). To test the effects of LOX inhibition on tumor initiation in vivo, we injected sgLOX-expressing A704 cells in mice and observed a significantly longer tumor-free survival compared to sgCntrl when using 40 mm^3^ as the predefined threshold for tumor initiation^45^ (**Fig. 4L**). Collectively, these findings demonstrate that LOX-HIF-2α axis drives ccRCC initiation and progression by regulating tumor cell plasticity and metastasis.

### Inhibition of LOX in tumor or endothelial cells impairs angiogenesis in vitro, ex vivo, and in vivo

Angiogenesis is a defining characteristic of ccRCC and is driven by HIF-2α-dependent transcriptional programs^46,47^ that we showed to be regulated by LOX. Therefore, we tested if LOX contributes to angiogenesis by performing complementary in vitro, ex vivo, and in vivo assays (**Fig. 5A**). In HUVEC tube formation assays, conditioned media from LOX-overexpressing HEK293T cells (LOX-CM), mimicking secreted LOX in TME and the angiogenic factors downstream of LOX-HIF-2α, significantly increased endothelial network complexity, junctions, nodes, and branching, at a level comparable to endothelial growth media (**Fig. 5B, C & Supplementary Fig. 6A, B**). Notably, scRNA-seq analysis of patient tumors showed that LOX is highly expressed in endothelial tip cells (**Fig. 5D**), a specialized migratory endothelial subset responsible for sprouting angiogenesis^48^. These LOX-expressing tip cells co-express canonical angiogenesis-associated genes, e.g., LY6H, implicating LOX in endothelial activation within the TME (**Fig. 5E**). We showed that LOX inhibition with LXG6403 in endothelial cells reduced tube formation to a degree similar to VEGFR blockade with axitinib, and their combination further suppressed tube metrics **(Fig. 5F, G, Supplementary Fig. 6C**). To specifically test the effects of inhibiting LOX in endothelial tip cells, we performed ex vivo aortic ring assay using aortic explants isolated from non-tumor-bearing C57BL/6 mice. This showed that LXG6403 treatment significantly decreases microvessel sprouting that is known to be regulated by endothelial tip cells, similar in magnitude to the reduction observed with axitinib treatment (**Fig. 5H, I**). These data confirm that inhibiting LOX (either in tumor cells or in endothelial cells) reduces angiogenic capacity in vitro and ex vivo.

**Figure 5.**
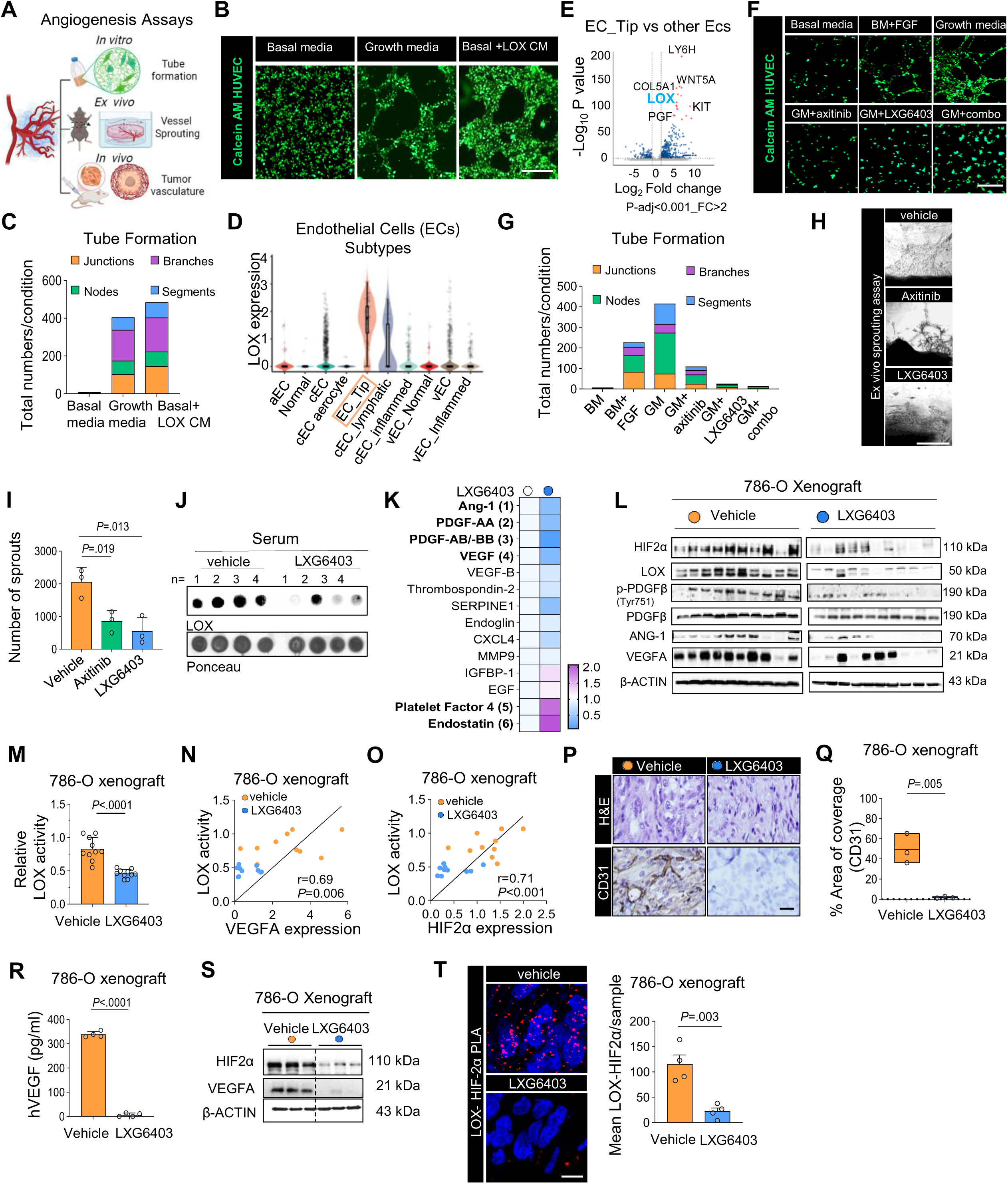
Inhibition of LOX in tumor or endothelial cells impairs angiogenesis in vitro, ex vivo, and in vivo. (**A**) Schematic overview of angiogenic assays performed. Created with BioRender.com. **(B)** Representative images of HUVEC tubule formation cultured with conditioned media from LOX-overexpressing HEK293T cells. HUVECs were labeled with calcein AM (n = 2). Scale bar = 200 μm **(C)** Quantitative analysis of the number of junctions, nodes, segments, and branches using Angiogenesis Analyzer (Image J) software (n = 2). **(D)** Single-cell RNA-seq analysis of patient tumors showing enriched LOX expression in endothelial tip cells, a subset critical for sprouting angiogenesis (n = 14 patient samples). (EC: endothelial cells; aEC: arterial EC; vEC: venous EC; cEC: capillary EC). (**E)** Volcano plot of differentially expressed genes in tip vs. non-tip endothelial cells, highlighting angiogenesis-associated transcripts. Differentially expressed genes (blue) are highlighted based on statistical significance and fold-change, the top 17 upregulated genes (red) are shown with gene labels. **(F)** Representative images of HUVEC tubule formation under growth factor stimulation, LXG6403, axitinib, or combination; HUVECs were labeled with calcein AM. Images are representative of (n = 3). Scale bar = 200 μm. **(G)** Quantitative analysis of the number of junctions, nodes, segments, and branches was done as in (f) (n = 2). **(H)** Representative images of ex vivo aortic ring assay showing microvessel sprouting from aortic explants isolated from non-tumor-bearing C57BL/6 mice treated with vehicle, LXG6403 (75 mg/kg, BID, p.o), and axitinib (30 mg/kg, BID, p.o) for 3 days. Each aortic explant was derived from a biologically independent mouse (n = 3 female C57BL/6 mice). Scale bar = 200μm **(I)** Quantification of (H), (n = 3 mice). **(J)** Serum LOX levels in NSG mice treated with vehicle and LXG6403 (150 mg/kg, p.o, 8h) (n = 4 mice per group). **(K)** Angiogenic protein array from sera collected from tumor-bearing mice following LOX inhibition, showing suppression of pro-angiogenic factors (VEGFA, PDGF-AA/AB, ANG-1) and induction endostatin, PF4 after LOX inhibition (n = 2 mice). **(L)** Immunoblot analysis of HIF-2α, LOX, p-PDGFβ, ANG-1 and VEGFA in tumors from 786-O xenografts treated for 8 h with vehicle and LXG6403 (100 mg/kg, p.o) (n = 10 mice). **(M)** Relative LOX activity in tumors from (L) (n = 10 mice). **(N, O)** Correlation analysis of tumor LOX activity with VEGFA **(N)** and HIF-2α **(O)** protein levels from n = 10. **(P)** Representative H&E and CD31 IHC images showing reduced vessel density with LXG6403 (n = 4 mice). Scale bar = 50μm **(Q)** Quantification of CD31+ area (% coverage) in tumors from (P) (n = 3 mice). **(R)** Human VEGFA (hVEGFA) serum ELISA showing reduced tumor-derived VEGFA with LXG6403 (n = 4 mice). **(S)** Immunoblot analysis of 786-O tumor lysates from (P) showing decreased HIF-2α and VEGFA levels after treatment (n = 3 tumors). **(T)** Proximity ligation assay (PLA) in tumor tissues showing reduced LOX-HIF-2α interaction following long-term LOX inhibition (n = 4 tumors). Scale bar = 25μm. Data are shown as mean ± SD for biological experiments, unless otherwise indicated. In vivo data are shown as mean ± SEM. Statistical significance was determined using unpaired two-tailed Student’s *t*-test (I, M, Q, R, T) or Pearson’s correlation (N, O). *P*-values are indicated.

To test the effects of LOX inhibition on angiogenesis in vivo, we treated 786-O xenografts with a single dose of LXG6403 for 8 hours, which resulted in a significant decrease in systemic LOX levels (**Fig. 5J, Supplementary Fig. 6D**), without histopathological changes in normal tissues (**Supplementary Fig. Fig. 6E**). Notably, angiogenic protein array analysis revealed that LXG6403 treatment broadly suppressed multiple pro-angiogenic factors, including VEGF, PDGF-AA/AB, and ANG-1 while increasing endostatin and platelet factor 4 levels (**Fig. 5K, Supplementary Fig. 6F**). Consistently, tumor lysates showed a reduction in p-PDGFRβ, ANG-1, and VEGFA levels upon LOX inhibition (**Fig. 5L**).

**Figure 6.**
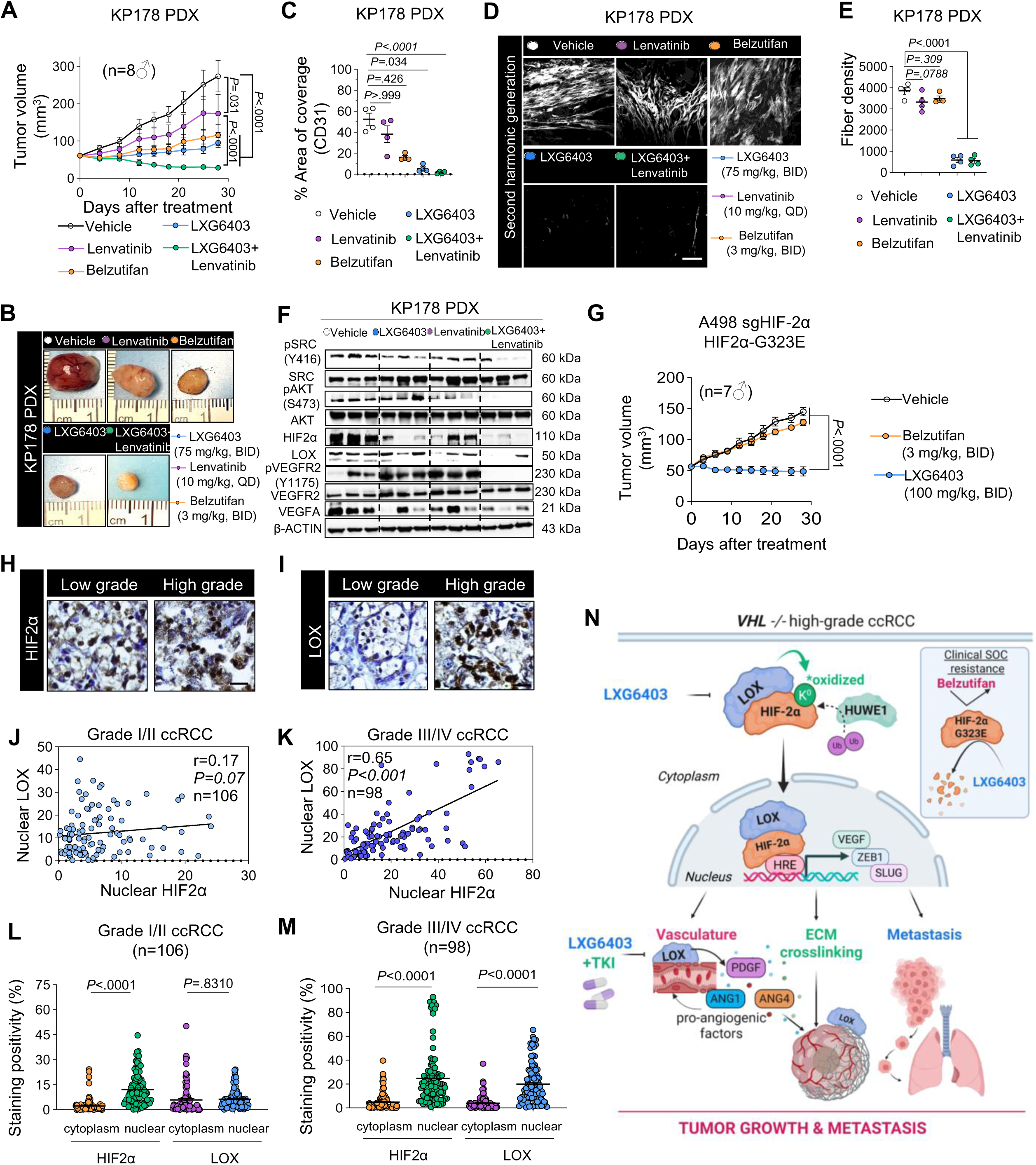
Targeting LOX enhances anti-angiogenic therapy and bypasses resistance to HIF-2α antagonism, and LOX and HIF-2α protein expressions correlate in high-grade ccRCC. (A, **B)** Tumor volume curves (A), and representative tumor images (B) of KP178 patient-derived xenograft (PDX) treated with vehicle, lenvatinib (10 mg/kg, QD), LXG6403 (75 mg/kg, BID), or combination (LXG6403 + lenvatinib) (n = 8 male NSG mice per group). **(C)** Quantification of CD31-positive area in tumor sections from (A) (n = 8 mice per group). **(D)** Representative second harmonic generation (SHG) imaging showing reduced collagen organization following LOX inhibition in KP178 ccRCC PDX tumors from (A) (n = 4 tumors). **(E)** Quantification of collagen fiber density, confirming disruption of the ECM structure upon LOX inhibition (n = 4 tumors). **(F)** Immunoblot showing reduced HIF-2α, LOX, p-SRC, p-AKT, p-VEGFR2, and VEGFA in tumors from KP178 PDX (A) (n = 3 mice). **(G)** Tumor volume curves of xenografts generated from A498 sgHIF-2α cells reconstituted with the resistance-associated HIF-2α G323E mutant and treated with vehicle, belzutifan (3 mg/kg, BID, p.o.), or LXG6403 (100 mg/kg, BID, p.o.) for 28 days (n = 7 male mice per group). **(H, I)** Representative TMA IHC images of HIF-2α (h) and LOX (I). Scale bar = 40 µm (n = 206 ccRCC patient tumors). **(J)** Correlation analysis between nuclear HIF-2α and nuclear LOX expression in low-grade (Grade I/II) ccRCC tumors (n = 106 patient tumors). **(K)** Correlation analysis between nuclear HIF-2α and nuclear LOX expression in high-grade ccRCC tumors (n = 100 patient tumors). **(L)** Quantification of nuclear vs cytoplasmic HIF-2α and LOX staining positivity in low grade (Grade I/II) ccRCC tumors. (n = 106 patient tumors). **(M)** Quantification of nuclear vs cytoplasmic HIF-2α and LOX staining positivity in high grade (Grade III/IV) ccRCC tumors. (n = 100 patient tumors). **(N)** Proposed model. Created with BioRender.com. Data are shown as mean ± SEM, unless otherwise indicated. Statistical significance was determined using one-way ANOVA with Dunnett’s (C, E), Holm’s (A, G), and Sidak’s (L, M) multiple-comparison corrections or Pearson correlation analysis (J, K). *P*-values are indicated.

Since HIF-2α transcriptionally regulates VEGFA, and LOX modulates HIF-2α stability and transcriptional activity, we next analyzed the LOX-HIF-2α-VEGFA axis in vivo. Short-term LOX inhibition (150 mg/kg LXG6403, P.O. for 8 hours) decreased both LOX enzymatic activity and HIF-2α-VEGFA levels (**Fig. 5L, M**). Importantly, we found that LOX activity positively correlates with both VEGFA and HIF-2α levels in tumors (**Fig. 5N, O**). Furthermore, we showed reduced microvascular density upon long-term LOX inhibition (**Fig. 3O**) as assessed by CD31 staining (**Fig. 5P, Q**), a decrease in human-specific circulating VEGFA (**Fig. 5R**), and reduced HIF-2α and VEGFA levels (**Fig. 5S**). Consistent with these findings, PLA in tumor tissues showed a marked reduction in LOX-HIF-2α interaction following long-term LOX inhibition (**Fig. 5T**), supporting disruption of this regulatory axis in vivo. Together, these data demonstrate that LOX sustains angiogenesis through regulation of HIF-2α-driven transcriptional programs, and that LOX inhibition suppresses hypoxia-driven angiogenic signaling and vascular organization.

### LOX inhibition enhances anti-angiogenic therapy response and remains effective in tumors harboring the belzutifan-resistant HIF-2α G323E mutation

We next tested the effects of LOX inhibition on enhancing response to anti-angiogenic therapy in vivo using a clinically relevant, highly angiogenic KP178 ccRCC PDX model. While lenvatinib monotherapy had a limited impact on tumor growth, LOX inhibition stabilized tumor growth (**Fig. 6A, B**). Remarkably, the combination of LXG6403 with lenvatinib yielded significant tumor regression (**Fig. 6A, B**). Importantly, treatment was well tolerated with no significant changes in body weight (**Supplementary Fig. 7A**), or hematologic parameters (e.g., RBC, HGB, HCT, WBC, platelets) observed across treatment groups (**Supplementary Fig. 7B**). Furthermore, CD31-positive vascular structures were markedly reduced upon combination treatment, together with increased apoptosis, assessed by TUNEL staining (**Fig. 6B, C, Supplementary Fig. 7C, D**). In addition, second harmonic generation (SHG) imaging showed that LOX inhibition markedly disrupted collagen organization (**Fig. 6D, E**), indicating that LOX uniquely regulates collagen organization and ECM architecture within the tumor microenvironment. Notably, p-VEGFR2 and downstream p-SRC, p-AKT levels were significantly reduced upon lenvatinib and LXG6403 combination treatment (**Fig. 6F)**. In the same experiment, HIF-2α inhibitor belzutifan also suppressed tumor growth (**Fig. 6A, B)**, with an effect comparable to LXG6403, and reduced CD31-positive vascular structures (**Fig. 6C)** without affecting collagen organization (**Fig. 6D, E)**.

Given the increasing clinical use of HIF-2α inhibitors and the emergence of resistance, associated with PAS-B pocket mutations in patients with advanced ccRCC^26^, we next tested whether LOX inhibition remains effective in tumors harboring the belzutifan-resistant HIF-2α G323E mutation. Given the increasing clinical use of HIF-2α inhibitors and the emergence of resistance, associated with PAS-B pocket mutations in patients with advanced ccRCC^26^, we next tested whether LOX inhibition remains effective in tumors harboring the belzutifan-resistant HIF-2α G323E mutation. Using HIF-2α-deficient ccRCC cells (sgHIF-2α), reconstituted with either wild-type HIF-2α or the resistance-conferring G323E mutant that we generated to model belzutifan resistance, we performed clonogenic assays upon treatment with belzutifan or LXG6403. As expected, belzutifan effectively suppressed colony formation in HIF-2α wild-type cells but failed to inhibit growth in cells expressing HIF-2α G323E. In contrast, LOX inhibition markedly reduced clonogenic growth in both wild-type and G323E-expressing cells (**Supplementary Fig. 7E-G**). Consistent with this, xenograft tumors generated using cells expressing HIF-2α G323E did not respond to belzutifan treatment, whereas they responded to LOX inhibition in vivo (**Fig. 6G**), with no significant changes in body weight (**Supplementary Fig. 7H**). Altogether, these results show that LOX inhibition enhances anti-angiogenic therapy response and remains effective in tumors harboring the belzutifan-resistant HIF-2α G323E mutation.

### LOX is associated with high-grade ccRCC and correlates with HIF-2α protein expression

To assess the clinical relevance of LOX-HIF-2α signaling in ccRCC, we analyzed LOX and HIF-2α protein expression and subcellular localization in ccRCC patient tumors (n = 206) (**Fig. 6 H, I**). High grade (Grade III/IV) tumors showed a significant positive correlation between nuclear LOX and nuclear HIF-2α levels, whereas there was no significant correlation in low grade (Grade I/II) tumors (**Fig. 6J, K**), suggesting a potentially more important role for LOX in more aggressive disease. Consistent with this, quantitative subcellular analysis revealed a progressive, grade-dependent increase in nuclear LOX localization, while nuclear HIF-2α levels were broadly elevated across tumor grades (**Fig. 6L, M**). Together, these data highlight a clinically relevant LOX-HIF-2α axis in which nuclear accumulation of LOX emerges as a potential defining feature of advanced disease, linking sustained HIF-2α-driven transcription program to tumor aggressiveness and therapeutic resistance in ccRCC.

## Discussion

While genomic studies have defined the mutational landscape of ccRCC, and bulk transcriptomic analyses have identified several prognostic genes^49–51^, little is known about the molecular drivers that sustain aggressive transcriptional states within malignant cell subpopulations. Here, we uncovered a new role for LOX, canonically known to remodel ECM, in sustaining a specific EMT-related gene program in malignant cells through stabilizing HIF-2α, thereby promoting EMT, angiogenesis, and disease progression. Our findings stem from foundational observations on RCC single-cell gene programs, demonstrating that malignant ccRCC cells organize into distinct transcriptional programs rather than uniform cellular states, revealing extensive intratumoral heterogeneity at the single-cell level^31^. Among these programs, GP60 emerged as a clinically relevant hypoxia-EMT-associated malignant program selectively active in ccRCC and associated with worse survival. *SERPINE1* and *LOX* were among the top-ranked genes associated with hypoxia and EMT signatures within the GP60 module. SERPINE1 has previously been implicated in ccRCC progression^32^. SERPINE2, another member of the serpin superfamily, was shown to promote RCC metastasis^52^, further supporting the robustness of our computational approach to elucidate ccRCC transcriptional diversity and pinpoint clinically-relevant gene programs. Although our data define LOX as a critical regulator of HIF-2α stability, nuclear accumulation, and chromatin occupancy, the precise genome-wide consequences of LOX-mediated regulation of HIF-2α transcriptional programs remain to be fully defined. Future studies using higher-resolution chromatin profiling will be required to determine how LOX selectively reshapes HIF-2α chromatin binding landscapes and downstream transcriptional output. ccRCC is defined by constitutive pseudohypoxia, pronounced intratumoral heterogeneity, and extreme hypervascularity, features that together promote tumor progression and limit therapeutic efficacy. Here, we identify LOX as a key driver of ccRCC progression and a therapeutically relevant target. We show, for the first time, that LOX stabilizes HIF-2α through oxidation-mediated blockade of HUWE1-dependent ubiquitination, thereby preventing proteasomal degradation and sustaining HIF-2α-driven transcriptional programs in ccRCC. LOX enhances HIF-2α chromatin occupancy in the nucleus, a finding clinically supported by the strong correlation between nuclear LOX and HIF-2α levels in high-grade ccRCC tumors. LOX sustains HIF-2α-driven transcriptional programs to promote tumor growth, plasticity, EMT, and angiogenesis. In parallel, LOX remodels the ECM through collagen crosslinking within the tumor microenvironment. Thus, LOX coordinates intracellular HIF-2α signaling with ECM remodeling and tumor vascularization in ccRCC. Inhibition of LOX promotes HUWE1-mediated HIF-2α degradation and disrupts ECM organization, resulting in suppression of tumor growth. Furthermore, targeting LOX hinders cell plasticity and tumor-initiating capacity, leading to reduced metastatic potential. Importantly, targeting LOX enhances sensitivity to anti-angiogenic therapy and remains effective in models harboring clinically relevant HIF-2α PAS-B pocket mutations (**Fig. 6N**). Overall, targeting LOX disrupts tumor-intrinsic (hypoxia signaling) and tumor-extrinsic (ECM remodeling and angiogenesis) processes simultaneously.

Oxygen-dependent prolyl hydroxylation and VHL-mediated ubiquitination have long been considered to be the primary mechanisms regulating HIF-2α protein stability^53–55^. However, three critical paradoxes still exist: (1) why HIF-2α levels and activity vary substantially among VHL-deficient tumors; (2) how high HIF-2α signaling is maintained in advanced disease, and (3) what mechanisms amplify HIF-2α transcriptional output beyond *VHL* loss. We show, for the first time, that LOX provides an additional layer of post-translational control by directly interacting with HIF-2α and oxidizing specific lysine residues; thereby impairing HUWE1-dependent ubiquitination and limiting proteasomal degradation. Furthermore, we show that LOX not only stabilizes HIF-2α, but it also acts as its cofactor, promoting its chromatin occupancy and transcriptional activity. These data were further supported by the strong positive correlation that we observed between nuclear LOX and HIF-2α at the protein level, specifically in high-grade ccRCC tumors, as well as in a grade-dependent increase in nuclear LOX localization. Together, these findings support the upstream regulatory role of LOX in reinforcing HIF-2α-associated transcriptional programs that promote ccRCC progression, uncovering a new mechanistic explanation for how HIF-2α levels are sustained in advanced disease.

Beyond novel biological insights, our findings have significant clinical implications. We show that LOX contributes to angiogenesis in ccRCC through HIF-2α-associated pro-angiogenic signaling and ECM remodeling. LOX inhibition reduced HIF-2α and VEGFA expression, impaired endothelial sprouting, reduced tumor vascularization, and disrupted fibrillar collagen organization in vivo. These findings identify a previously unrecognized role of LOX in functionally linking hypoxia signaling with ECM remodeling during angiogenesis. Clinically, angiogenesis is a major therapeutic dependency in ccRCC due to constitutive HIF activation and VEGF-driven hypervasculature following VHL loss^56–58^. Consequently, VEGFR-targeted therapies remain a cornerstone of treatment for advanced ccRCC, yet responses are often limited by persistent hypoxia signaling and treatment-associated toxicities. Therapeutic benefit from LOX inhibition has the potential to go beyond VEGFR blockade by simultaneously reducing HIF-2α stability and disrupting ECM organization.

HIF-2α inhibitors^19,23,24^ represent one of the major therapeutic advancements in ccRCC and demonstrated meaningful clinical benefit by suppressing HIF-2α-dependent transcriptional activity^59^.^59^. However, several clinically relevant limitations have emerged. Although these agents hinder transcriptional activity of HIF-2α, they do not inhibit the upstream mechanisms that regulate HIF-2α protein stability and degradation in advanced disease. As a result, on-target systemic toxicities such as anemia observed in clinical trials are inevitable, because HIF-2α is also expressed in normal cell populations, including erythroid lineages^60^. Furthermore, acquired PAS-B domain mutations can disrupt inhibitor binding, limiting the durability of the response^25,26^.As a result, on-target systemic toxicities such as anemia observed in clinical trials are inevitable, because HIF-2α is also expressed in normal cell populations, including erythroid lineages^60^. Furthermore, acquired PAS-B domain mutations can disrupt inhibitor binding, limiting the durability of the response^25,26^. Here, we show that unlike HIF-2α inhibitors, LOX targeting promotes HIF-2α protein degradation, reducing the available HIF-2α pool independently of PAS-B domain integrity and remaining effective despite resistance-conferring PAS-B mutations, without any observable toxicity.

Together, our findings establish LOX as a central regulator of hypoxia signaling and tumor progression in ccRCC. We propose that targeting LOX offers a multi-pronged therapeutic strategy to destabilize HIF-2α, reprogram the tumor microenvironment, and address resistance in advanced ccRCC, with potential applicability across other hypoxia-driven solid malignancies.

## Methods

### Patient Samples and Ethical Approval

Primary and metastatic RCC specimens used for single-cell transcriptomic analyses were obtained from patients undergoing nephrectomy following written informed consent under protocols approved by the McGill University Health Care Research Ethics Board (MUHC REB), as described in Madrigal et al^31^.

For the IHC studies, ccRCC tissue microarrays (TMAs) containing primary tumor specimens were obtained from the McGill University Health Centre under protocols approved by MUHC REB following written informed consent. Clinicopathologic characteristics of the cohort included in the present study are summarized in **Supplementary Table 1**. Histologic grade was available for all specimens, whereas age, sex, and pathologic T stage were available for a subset of patients. All samples were reviewed by board-certified genitourinary pathologist to confirm diagnosis and histopathological annotations. The remaining clinicopathological characteristics of the cohort have been described previously^61^. TMA sections were used for the IHC analyses of LOX and HIF-2α. For the IHC studies, ccRCC tissue microarrays (TMAs) containing primary tumor specimens were obtained from the McGill University Health Centre under protocols approved by MUHC REB following written informed consent. The clinicopathological characteristics of this cohort have been described previously^61^. All samples were reviewed by board-certified genitourinary pathologist to confirm diagnosis and histopathological annotations. TMA sections were used for the IHC analyses of LOX and HIF-2α.

### Single-cell RNA Sequencing and Analysis

Processed single-cell expression data, including cell type annotations and malignant cell identification, were obtained from^31^. Gene expression programs identified using single-cell convex non-matrix factorization were also retrieved. Gene set activity was estimated using GEDI (v.1.0.2), incorporating gene sets from the Hallmark MSigDB collection, a HIF2α signature, and HIF-derived gene signatures^31^. Dimensionality reduction was performed using Uniform Manifold Approximation and Projection (UMAP), implemented in the uwot package (v.0.1.16). Survival analysis of gene program activity was conducted as described^31^. Differentially expressed genes in the tip-cells (cEC_LOX) over other endothelial cells (ECs) were identified using a Wilcoxon Rank Sum test, and *P*-values were adjusted by Bonferroni correction using all genes in the data. Differentially expressed genes (p-adj < 0.001 & FC > 2) are colored with blue while the top 20 upregulated genes are highlighted as red color with gene labels.

### Clinical and Transcriptomic Data Analysis

Gene expression and clinical outcome data for ccRCC patients were obtained from The Cancer Genome Atlas Kidney Renal Clear Cell Carcinoma (TCGA-KIRC) cohort and Gene Expression Omnibus (GEO) datasets (GSE40435, GSE152938). Additional scRNA-seq data of normal kidneys (n = 20) as well as major RCC subtypes were processed and analyzed together with our scRNA-seq data (GSE202109, GSE131685). Expression of LOX was visualized by log *P* transformed normalized expression (proximal (n = 8263), non-proximal (n = 1879), (ccRCC n = 19470)). Expression of E3 ubiquitin ligases in ccRCC tumors vs. normal in TCGA was downloaded from the UCSC Xena browser (https://xenabrowser.net/.) Pearson correlation analyses between LOX expression and other genes of interest were performed. For survival analyses, patients were stratified into high and low GP60 activity or LOX expression groups based on median expression values, and analyses were performed using Cox proportional hazards models, with Kaplan-Meier estimates compared using log-rank tests. Where applicable, *P*-values were adjusted for multiple testing using the Benjamini-Hochberg method. All analyses were performed using GraphPad Prism (v10.0) and R statistical software.

### Exonic and Intronic Read Analysis

Exonic and intronic read counts were quantified using velocyto (v.0.17.17). For each sample, exonic and intronic counts were normalized separately using the logNormCounts function from the scuttle (v.1.12.0) package. Pearson correlation analyses were then performed between log-normalized LOX intronic and exonic expression and selected gene sets.

### Cell Culture

ccRCC cell lines (786-O; RRID: CVCL_1051, A704; RRID: CVCL_1065, A498; RRID: CVCL_1056, CAKI-1; RRID: CVCL_0234, and ACHN; RRID: CVCL_1067; ATCC), and their LOX-knockdown derivatives were cultured in RPMI-1640 medium (Corning, Cat. no.10041) supplemented with 10% fetal bovine serum (Gibco, Cat. no.16140071) and 50 U/ml penicillin/streptomycin (Corning, Cat. no.15140122) at 37°C in a humidified atmosphere containing 5% CO_2_. HEK293T cells were maintained in Dulbecco’s modified Eagle medium (Gibco, Cat. no.10014) with 10% fetal bovine serum (Gibco, Cat. no.16140071) and 50 U/ml penicillin/streptomycin (Corning, Cat. no.15140122) under identical conditions. Cells were used within 10-20 passages from the initial thawing. Human umbilical vein endothelial cells (HUVECs) were maintained in endothelial growth medium (EGM-2; Lonza, Cat. no.CC3162) and used within passages 2-5. All cell lines were authenticated by short tandem repeat profiling and confirmed mycoplasma-free by PCR testing using Lookout One-Step mycoplasma detection kit (Sigma, Cat. no.MP0035).

### Genetical and pharmacological perturbations

LOX knockouts were generated using lentiviral CRISPR/Cas9 constructs containing two independent sgRNAs targeting LOX (ABM Good, Cat. no.275401110595). LOX inducible knockdown models were generated as previously described^41^. Briefly, TRIPZ inducible lentiviral non-silencing shRNA Control (RHS4743) and LOX shRNA vectors were obtained from Dharmacon. To produce viral particles, 6 μg of each of these vectors and 4.3 μl of trans-lentiviral packaging mix were used to co-transfect HEK293FT cells in 6-wells plate with CaCl2 reagent. 48 hours post-transfection, viral particles were collected and used to transduce 786O cells. Stable cell lines were selected using puromycin (1 μg/mL) for 3-4 days following transduction. LOX inhibitor LXG6403 was synthesized as reported previously^42^. Belzutifan (Cat. no. HY-125840), axitinib (Cat. no. HY-10065), lenvatinib (Cat. no. HY-10981), BI8622 (HUWE1 inhibitor, Cat. no. HY-120929), and heclin (Cat. no. HY-110204) were purchased from MedChemExpress.

### Protein stability and ubiquitination assays

For half-life assays, cells were pre-treated with cycloheximide (100 µg/ml, Sigma-Aldrich) and harvested at the indicated time points for immunoblot analysis. For ubiquitination assays, cells were pre-treated with MG132 (10 µM, 6 h; MedChemExpress, Cat. no.13259), lysed under denaturing conditions (1% SDS, 50 mM Tris-HCl pH 7.5, 150 mM NaCl, 10 mM N-ethylmaleimide, protease inhibitors), and diluted 10-fold before immunoprecipitation with anti-FLAG antibody overnight at 4°C (**Supplementary Table 2)**. Immunoprecipitates were resolved by SDS-PAGE and immunoblotted with anti-Ubiquitin and anti-HIF-2α or anti-FLAG antibodies **(Supplementary Table 2)**.

### In vitro oxidation and ubiquitination assays

For the detection of LOX-mediated oxidative modifications, protein carbonylation was assessed using biotin hydrazide labeling. FLAG-tagged HIF-2α was immunopurified from HEK293T cells using anti-FLAG magnetic beads (Thermo Fisher Scientific, Cat. no.PIA36797) under native conditions. Bead-bound HIF-2α was incubated with recombinant human LOX (Origene, Cat. no.TP313323) in oxidation buffer (50 mM HEPES and 10 mM CaCl_2_, pH 8.0) at 37 °C for 45 min in the presence or absence of LXG6403. After incubation, the beads were washed to remove the LOX enzyme. Following oxidation, carbonyl groups were labeled by incubating bead-bound proteins with EZ-Link Biotin Hydrazide (Thermo Fisher Scientific, Cat. no.21339) for 2 h at room temperature with rotation. Beads were washed extensively with TBS-T, and proteins were eluted in SDS sample buffer, resolved by SDS-PAGE, and transferred to PVDF membranes. Biotin-labeled carbonylated proteins were detected by immunoblotting using streptavidin-HRP followed by chemiluminescent detection.

For in vitro ubiquitination assays, bead-bound HIF-2α (oxidized or control) was incubated with ubiquitination reaction components (E1, E2, ubiquitin, ATP; Enzo Life Sciences, Cat BML-UW9920-0001) supplemented with HUWE1-IP eluted from 786-O cells as a source of E3 ligase activity. Samples were treated with the LOX inhibitor LXG6403 or the HUWE1 inhibitor BI8622, and reactions were carried out at 37°C for 60 min and terminated by the addition of SDS sample buffer. Ubiquitination was assessed by immunoblotting using streptavidin-HRP and anti-FLAG antibodies (**Supplementary Table 2**).

### Animal Experiments

All the in vivo studies were carried out in accordance with the Institutional Animal Care and Use Committee of the Medical University of South Carolina. Sex-balanced cohorts of NOD-scid IL2Rγ^null^ (NSG) mice (6-8 weeks old; The Jackson Laboratory; RRID: IMSR_JAX:005557) were used for xenograft and PDX studies whenever possible. Mice were subcutaneously injected with 10×10^6^ 786-O, 5×10^6^ A498, or 1×10^6^ A704 cells or implanted with patient-derived xenograft (PDX) tumor fragments (KP178) in the right flank. Mice were randomized into treatment groups once tumors reached comparable sizes. In the A704 xenograft metastasis experiment, the starting tumor volume was 75 mm³, and the treatments were vehicle, belzutifan (3 mg/kg, BID), and LXG6403 (75 mg/kg, BID). In the KP178 ccRCC PDX experiment, the starting tumor volume was 60 mm³, and mice were treated with vehicle, lenvatinib (10 mg/kg, QD), belzutifan (3 mg/kg, BID), LXG6403 (75 mg/kg, BID), and combination therapy (LXG6403 plus lenvatinib). In single-agent experiments, starting tumor volume was 50 mm³ for the 786-O xenograft model, and treatment used LXG6403 (100 mg/kg, BID). For HIF-2α-G323E xenograft studies, mice bearing tumors derived from wild-type or G323E HIF-2α-reconstituted cells were treated with vehicle, belzutifan (3 mg/kg, BID) or LXG6403 (100 mg/kg, BID). All treatments were administered by oral gavage (p.o.) for 21-28 days.

Tumor volumes were measured twice weekly using digital calipers and calculated using the formula: volume = (length × width^2^)/2. Body weights were recorded twice weekly to monitor toxicity. At the experimental endpoint, mice were euthanized, and tumors were harvested, weighed, and processed for histological analysis, protein extraction, and ELISA. Tissue sections were stained with hematoxylin and eosin (H&E), terminal deoxynucleotidyl transferase dUTP nick end labeling (TUNEL staining; apoptosis).

### Immunohistochemistry and Imaging

Formalin-fixed, paraffin-embedded (FFPE) ccRCC tumor sections were processed for immunohistochemistry using standard deparaffinization, rehydration, and heat-induced antigen retrieval protocols. Endogenous peroxidase activity was quenched, and non-specific binding was blocked before incubation with primary antibodies against LOX (1:600) or HIF-2α (1:600) overnight at 4°C. Signal detection was performed using a biotin-streptavidin horseradish peroxidase system with 3,3′-diaminobenzidine (DAB) as the chromogen, followed by hematoxylin counterstaining. All staining was performed manually using the Abcam HRP/DAB detection IHC kit (Abcam, Cat. no. ab62264). Whole slides were imaged at 20x magnification using the PhenoImager® HT™ 2.0 Automated Quantitative Pathology Imaging System (Akoya Biosciences). Tumor regions were annotated using Phenochart software. Quantitative image analysis was performed using inForm® Tissue Analysis Software, for which a supervised machine-learning classifier was trained on representative tumor regions to distinguish nuclear and cytoplasmic compartments and to segment individual cells. Following classifier training and validation, nuclear and cytoplasmic scores were quantified. Spatial and correlative analyses were conducted using the PhenoptrReports open-source R package and GraphPad Prism (v10.0).

### Statistical Analysis

Statistical analyses were performed using GraphPad Prism (v10.0) and R statistical software (v4.3.0). Data are shown as mean ± standard deviation (SD) or mean ± standard error of the mean (SEM) as indicated in figure legends. Statistical details, including sample size (n), statistical tests, and significance thresholds, are provided in the corresponding figure legends. Unless otherwise indicated, n represents biological replicates rather than technical replicates. For in vivo studies, n refers to individual animals, whereas for in vitro studies, n refers to independent biological experiments, unless otherwise indicated in the figure legend. All experiments were independently replicated at least two or three times, as indicated in the figure legends. Statistical significance was determined using unpaired two-tailed Student’s t-tests for comparisons between two groups or one-way analysis of variance (ANOVA) followed by Tukey’s multiple comparisons test for comparisons among multiple groups. For in vivo studies, linear mixed-effects (LME) models were fit using the nlme package (v3.1-164) with natural log-transformed tumor volumes as the outcome variable. The models included baseline tumor volume, time, treatment group, and time-treatment interaction as fixed effects, with random intercepts for individual mice. Body weight changes were modeled similarly using percent change from baseline. Linear contrasts were used to assess group differences at specific time points, with *P*-values adjusted for multiple comparisons using Holm’s method. Statistical tests were performed using emmeans package (v1.10.4) in R. For survival analyses, hazard ratios and 95% confidence intervals were calculated using Cox proportional hazards models. Sample sizes were estimated using power analyses (80% power and α = 0.05) to detect biologically meaningful differences.

All other methods are provided in Supplementary Methods.

### Data Availability

Single-cell RNA sequencing datasets analyzed in this study were obtained from^31^. This study also analyzes previously published, publicly available datasets. Accession numbers and references for these datasets are provided in the Clinical and Transcriptomic Data Analysis section.

## Acknowledgements

We thank the members of the Ozgur Sahin laboratory for their valuable discussions and advice. This work was supported by the United States Department of Defense (DoD KCRP) Idea Development Award W81XWH2110945 (Y.R. and O. Sahin), in part by the NIH (R01CA267101 and R01CA251374 to O. Sahin), and by the SmartState Endowment in Lipidomics and Drug Discovery (O. Sahin). The Biorepository and Tissue Analysis Shared Resource is supported in part by the Hollings Cancer Center, Medical University of South Carolina, through the NCI (P30 CA138313). The animal core facility utilized in this study was supported by the NIH (C06 RR015455). The Biostatistics Shared Resource at the Hollings Cancer Center, Medical University of South Carolina, was supported by the NIH (P30 CA138313). Computational resource allocations utilized were provided by the Digital Research Alliance of Canada to H.S.N. The Zeiss 880 microscope was funded by a Shared Instrumentation grant (S10 OD018113). H.S.N. holds a CIHR Canada Research Chair (CRC-2021-00423). A.M. and Z. M. were supported by doctoral training awards from FRQS.

## Author contributions

Conceptualization, B.U., A.M., M.K., H.N., Y.R. and O.S.; methodology, B.U., O.Saatci, A.M., M.K., W.T., M.S., Z.M., K.S., C.N.R., T.N., J.R., C.M., R.C.R., H.N., Y.R. and O.S.; investigation, B.U., O.Saatci, A.M., M.K., W.T., J.R., M.P., R.C.R., H.N., Y.R. and O.S.; in vivo experiments, B.U. and O.S.S.; formal analysis, B.U., O.Saatci, A.M., M.K. and W.T.; statistical analysis, J.S.A. and E.H.; writing original draft, B.U.; writing-review & editing, B.U., O.Saatci, A.M., M.K., W.T., J.R., C.M., M.P., F.B., S.T., R.C.R., H.N., Y.R. and O.S.; resources, M.S., Z.M., T.N., V.P., J.R., C.M., M.P., F.B., S.T., H.N., Y.R. and O.S.; funding acquisition, Y.R. and O.S.

## Supplementary Material

### SUPPLEMENTARY METHODS

#### Immunoblotting

Protein isolation and immunoblotting were performed as previously described^1^. Briefly, RIPA buffer was used to isolate total protein lysate in the presence of protease and phosphatase inhibitor cocktails, and protein concentrations were measured using the BCA Protein Assay Reagent Kit (Thermo Fisher Scientific, Cat. no. 23223, 23224). Equal amounts of protein were separated on 8-10% SDS-PAGE gel. Separated proteins were transferred onto PVDF membranes (Bio-Rad, Cat. no.1620177) using a Trans-Blot turbo transfer system (Bio-Rad) and incubated with primary antibodies at dilutions indicated in **Supplementary Table 2**. Horseradish peroxidase-conjugated anti-mouse or anti-rabbit antibodies (**Supplementary Table 2**) were used as secondary antibodies, and signals were detected by enhanced chemiluminescence (Thermo Fisher Scientific, Cat. no. A38556). Images were acquired using Image Lab Software (Biorad) or iBright Software (Invitrogen).

#### Quantitative RT-PCR (qRT-PCR) analysis

Total RNA was extracted from cultured cells using Quick-RNA Miniprep Kit (Zymo, Cat. no. R1055), and cDNAs were generated using RevertAid RT Reverse Transcription Kit (Thermo Fisher Scientific, Cat. no. K1622). qRT-PCR analysis was performed with gene-specific primers, using LightCycler 480 SYBR Green I Master kit (Roche). *HPRT1* and *ACTB* were used as housekeeping genes. The average Ct value was calculated from triplicates of each sample, and the relative mRNA expression was determined. Sequences of the qRT-PCR primers are listed in **Supplementary Table 3**.

#### Transient transfection with siRNAs and overexpression reporters

Transfections were carried out by seeding cells in media containing no antibiotics and changing media with Opti-MEM reduced serum medium (Thermo Fisher Scientific, Cat. no.31985070) prior to transfection with Lipofectamine 2000 (Thermo Fisher Scientific, Cat. no.11668019) in 6-well or 96-well format. siRNA transfections were performed at a final concentration of 100 nM (**Supplementary Table 4**), and RNA or protein was isolated after 48 hours of transfection. Overexpression plasmids were transfected at 500 ng per well in 6-well plates.

#### Co-immunoprecipitation

Cells were treated with LXG6403 (5, 15, and 30 µM) for 4 hours and lysed in lysis buffer (50 mM Tris-HCl pH 7.0, 150 mM NaCl, 0.2% NP40, 7.5% glycerol, protease and phosphatase inhibitor cocktails. Lysates were clarified by centrifugation, and 1 mg of total protein per condition was incubated overnight at +4 °C with Dynabeads Protein G (Thermo Fisher Scientific, Cat. no.10003D) pre-coated with the indicated antibodies (**Supplementary Table 2**). Beads were washed and resuspended in 60 µL 1X SDS sample buffer, boiled at 70 °C for 10 min, and loaded onto a polyacrylamide gel. Immunoblot was done using antibodies provided in **Supplementary Table 2**.

#### Chromatin immunoprecipitation (ChIP) assay

ChIP assays were performed as described previously^2^. Briefly, 786-O cells that were grown to 70% confluency were crosslinked with 1% formaldehyde for 10 min, followed by quenching with 125 mM glycine for 5 min. Cells were lysed in 500 µL lysis buffer, and nuclear lysates were extracted.

Sheared chromatin was incubated overnight at 4 °C with HIF-2α antibody and magnetic beads with slow agitation (**Supplementary Table 2**). IgG antibody was used as the negative control. Samples were washed with low and high salt wash buffers, and Proteinase K treatment was done for 2 h at 62 °C with shaking. Samples were incubated at 95 °C for 10 min and separated from the beads using a magnetic separator. DNA was isolated, and ChIP-qPCR was performed using primers targeting the VEGFA, SERPINE1, PTHrP, and CCND1 promoters. The results were normalized to the bead-only control and represented as fold enrichment. Sequences of the ChIP-qPCR primers are listed in **Supplementary Table 5**.

#### Lysyl oxidase activity

Lysyl oxidase activity assay within cells or within tumors was measured using the fluorometric Lysyl Oxidase Activity Assay Kit (Abcam, Cat. no. ab112139) according to the manufacturer’s instructions. Briefly, serially diluted replicates of tumor lysates or cell supernatants were incubated with reagent mix and subsequently measured in a fluorescence microplate reader at Ex/Em = 540/590 nm wavelength. Tumor lysates were prepared by sonicating tissue samples in extraction buffer (6 M urea, 10 mM Tris, pH 7.4, and protease inhibitors) followed by centrifugation.

#### Cell fractionation and chromatin extraction

Cytoplasmic and nuclear protein fractions were isolated using the Nuclear Extract Kit (Active Motif, Cat. no.54001) according to the manufacturer’s protocol. Cells were lysed in a hypotonic buffer to release cytoplasmic content, followed by high-salt extraction of nuclear proteins. Fraction purity was validated by immunoblotting with cytoplasmic marker α-tubulin and nuclear marker lamin B1 (**Supplementary Table 2**). Resulting fractions were followed by immunoprecipitation.

For chromatin fractionation, cells were lysed using buffer (10 mM Tris-HCl pH 7.5, 10 mM NaCl, 3 mM MgCl₂, 0.5% NP-40, with protease/phosphatase inhibitors) and incubated for 45 min on ice. Lysates were centrifuged at 16,000 × g for 15 min at 4°C to separate the cytosolic fraction (supernatant). The pellet was washed once with 200 µL wash buffer (10 mM Tris-HCl pH 7.5, 10 mM NaCl, 3 mM MgCl₂, 1 mM EDTA), centrifuged, then resuspended in 200 µL wash buffer and sonicated (3 × 10 sec pulses). This chromatin fraction was quantified by BCA and analyzed by Western blot, loading 2× protein amounts compared to cytosol. Histone H3 and α-tubulin were used as fraction controls.

#### Tubule Formation and ex vivo angiogenesis assays

HUVECs were labeled with 2 μM calcein-AM (Invitrogen) for 30 min at 37°C prior to cell plating. Labeled cells (2 × 10^4^) were seeded on growth factor-reduced Matrigel-coated 96-well plates and cultured in endothelial growth medium (Lonza, Cat. no.CC3162) containing axitinib (1 μM; MedChemExpress, Cat. no. HY-10065), the LOX inhibitor LXG6403 (5 μM), or conditioned media derived from LOX-modulated HEK293T cells. Tube formation was assessed after 16-18 h of incubation. Fluorescent images were captured using an inverted fluorescence microscope, and tube length, branching points, and network area were quantified using the Angiogenesis Analyzer plugin in ImageJ^3^.

Thoracic aortas were isolated from non-tumor-bearing C57BL/6 mice (The Jackson Laboratory; RRID:IMSR_JAX:000664) and sectioned into 1-mm pieces. Aortic explants were embedded in growth factor-reduced Matrigel (Corning, Cat. no.356231) and cultured in endothelial growth medium (Lonza, Cat. no.CC3162) containing vehicle, LXG6403, or axitinib for 3 days. Microvessel sprouting was imaged using brightfield microscopy, and sprout formation was quantified using ImageJ software.

#### Colony formation, sphere assays, and 3D cultures

For colony formation, cells (1,000 cells/well) were seeded in 12-well plates and treated with the indicated inhibitors for 14-21 days. Colonies were fixed, stained with 0.2% crystal violet, and quantified at the experiment endpoint. Spheroid assays were performed using ultra-low attachment plates in serum-free DMEM/F12 (Thermo Fisher Scientific, Cat. no.11330032) supplemented with B27 (Thermo Fisher Scientific, Cat. no.17504044) EGF (20 ng/ml, Peprotech, Cat. no.100-15), and bFGF (20 ng/ml, Peprotech, Cat. no.100-26, 100-19). Sphere size and number were quantified after 14-21 days. Collagen-embedded 3D culture assays were performed as previously described^4^. Briefly, neutralized collagen matrices were prepared using rat tail collagen I (Corning, Cat. no.354236) at a final concentration of 1 mg/mL with pH adjustment using 1 N NaOH. Cell pellets were resuspended in the collagen mixture prepared in PBS and seeded into 96-well plates. Collagen matrices were allowed to solidify for 1 h at room temperature prior to the addition of culture media. Embedded cells were subsequently cultured at 37 °C for the indicated time periods, and drug treatments were initiated 12 h after seeding. Cell viability was assessed using the CellTiter-Glo 3D assay (Promega, Cat. no. G9683) according to the manufacturer’s instructions.

#### Migration and invasion assays

Transwell migration and matrigel invasion assays were performed using 8-µm pore size inserts (Corning, Cat. no.3428). Cells were plated in serum-free medium, allowed to migrate for 24 h (migration) or 48 h (invasion), then fixed with 4% PFA, stained with 0.2% crystal violet, and quantified using Image J.

#### Angiogenesis protein array

Serum angiogenic profiling was performed using the Proteome Profiler Mouse Angiogenesis Array Kit (R&D Systems, Cat. no. ARY015) according to the manufacturer’s instructions. Briefly, sera collected from treated mice were incubated with nitrocellulose membranes pre-spotted with angiogenesis-related capture antibodies. Membranes were processed following the manufacturer’s protocol, and images were acquired using Image Lab Software (BioRad). Relative intensity was quantified by densitometric analysis using ImageJ software.

#### Oxyblot Analysis of recombinant HIF-2α oxidation

To independently validate direct LOX-mediated oxidation, protein carbonylation was assessed using the OxyBlot Protein Oxidation Detection Kit (Sigma-Aldrich, Cat. no. S7150) with bacterially expressed recombinant HIF-2α. Purified recombinant HIF-2α was incubated with recombinant human LOX (Origene, Cat. no.TP313323) in oxidation buffer at 37°C for 45 min in the presence or absence of LOX inhibitor. Proteins were derivatized with 2,4-dinitrophenylhydrazine (DNPH) according to the manufacturer’s instructions, resolved by SDS-PAGE, transferred to PVDF membranes, and carbonylated proteins were detected using anti-DNP antibody. This assay was used as an orthogonal validation of LOX-dependent oxidation in a cell-free system.

## SUPPLEMENTARY FIGURES

**Supplementary Figure 1.**
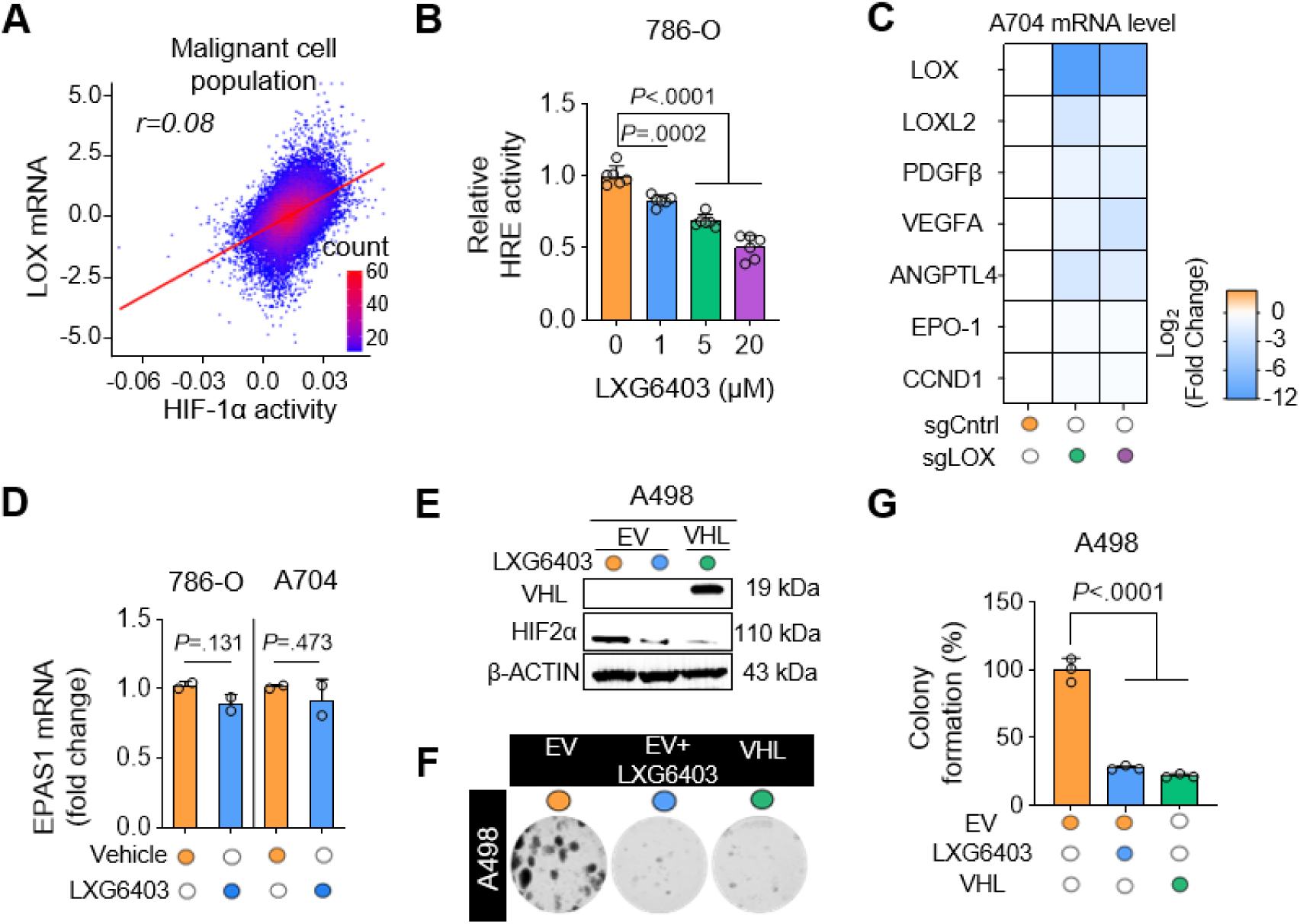
LOX inhibition reduces HIF-2α transcriptional activity without affecting *EPAS1* mRNA expression. **(A**) Relationship between LOX expression and HIF-1α activity in malignant RCC cells. Scatterplot showing latent-log expression of LOX (x-axis) versus HIF-1α activity (y-axis). The line represents a linear model fit, and Pearson correlation values are indicated. **(B)** HRE-GFP reporter assay showing dose-dependent reduction of HIF-2α transcriptional activity upon LOX inhibition in 786-O cells (n = 6 technical replicates). **(C)** qRT-PCR analysis of canonical HIF-2α targets (LOX, LOXL2, PDGFB, VEGF, ANGPTL4, EPO-1, CCND1) in A704 sgLOX cells (n = 2). **(D)** qRT-PCR analysis of *EPAS1* mRNA showing unchanged transcript levels following LXG6403 treatment, indicating post-translational regulation of HIF-2α (n = 2). **(E)** Western blot analysis of VHL and HIF-2α in A498 cells transfected with empty vector (EV) ± 7.5 µM LXG6403 or with wild-type VHL. β-ACTIN was used as a loading control here and in all immunoblots unless otherwise indicated. **(F)** Colony formation assay comparing A498 EV ± 7.5 µM LXG6403 and A498 cells expressing wild-type VHL. **(G)** Quantification of (F) (n = 3). Data are shown as mean ± SD. A two-sided Student’s t-test was used to calculate the statistical difference between the two groups (D). One-way ANOVA with Dunnett’s test was performed to compare more than one group (B, G). *P*-values are indicated. Colors denote experimental groups and are used consistently across graphs and representative images throughout the manuscript unless otherwise indicated. Open circles indicate control conditions, whereas filled circles indicate experimental conditions.

**Supplementary Figure 2.**
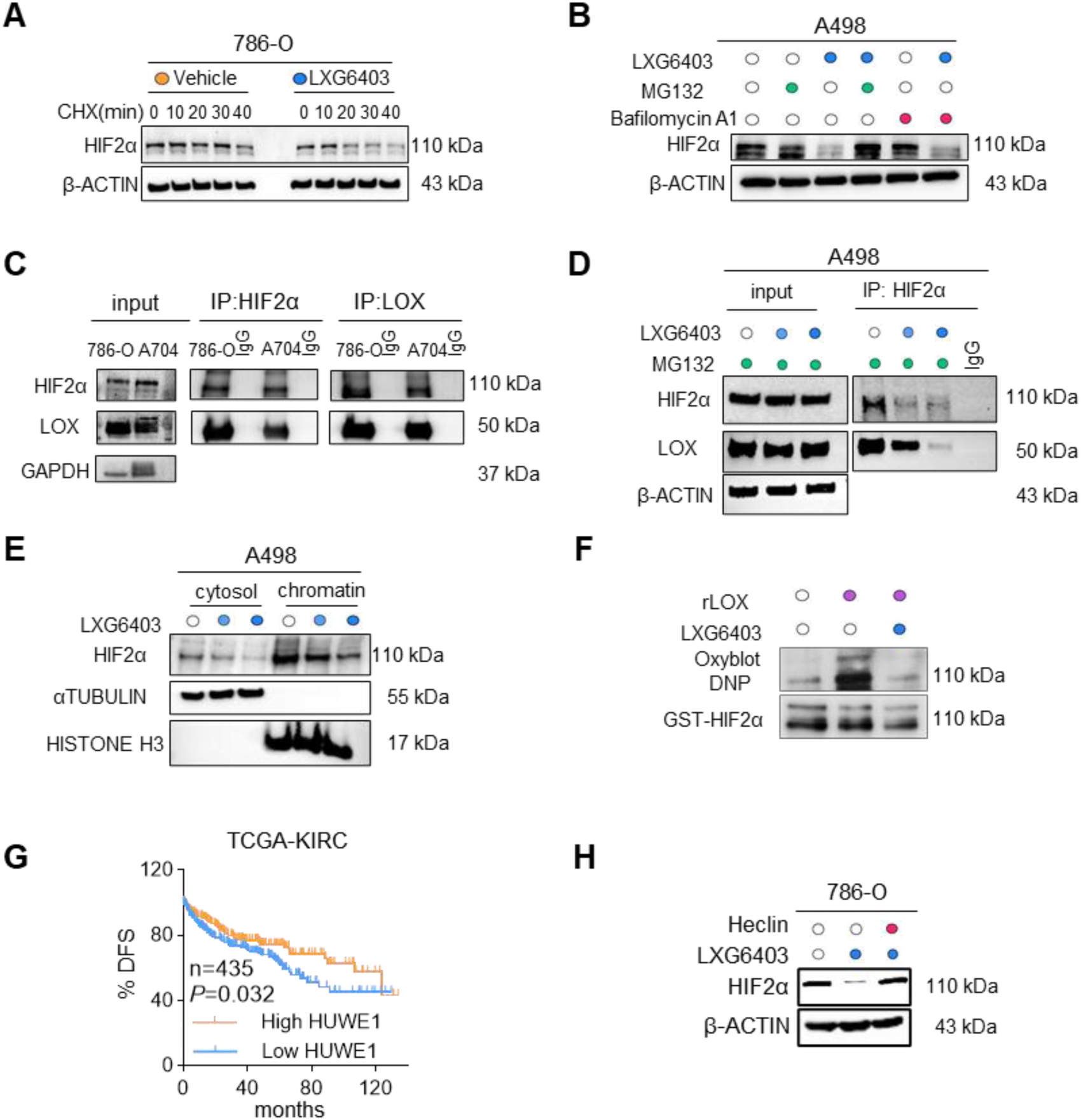
LOX stabilizes HIF-2α and LOX inhibition induces HUWE1-mediated proteasomal degradation of HIF-2α, resulting in loss of its chromatin occupancy. **(A)** Representative immunoblot from cycloheximide (CHX) chase assay showing accelerated HIF-2α degradation upon LOX inhibition. β-ACTIN was used as a loading control here and in all immunoblotting figures, unless otherwise indicated. **(B)** Western blot of 786-O cells treated with ± LXG6403 and ± MG132 or bafilomycin A1, showing that proteasome inhibition rescues HIF-2α abundance under LOX inhibition, whereas lysosomal inhibition does not (LXG6403 30µM for 24 hours, MG132 10µM for 6 hours, Bafilomycin A1 50 nM for 6 hours). **(C)** Co-immunoprecipitation (co-IP) and reverse-IP demonstrating a physical interaction between LOX and HIF-2α. GAPDH is used as a loading control. **(D)** Immunoblot of LOX and HIF-2α in A498 cells treated with LXG6403 for 24 hours in the presence of MG132 (10µM for the last 6 hours). Increasing blue color intensity corresponds to increasing LXG6403 concentrations (15 and 30 μM). **(E)** Chromatin fractionation assays showing reduced chromatin-bound HIF-2α upon LOX inhibition in a dose-dependent manner. Increasing blue color intensity corresponds to increasing LXG6403 concentrations (15 and 30 μM) for 48 hours. α-tubulin and Lamin B1 serve as fractionation control. **(F)** OxyBlot analysis of a bacterial cell-free system demonstrating LOX-mediated oxidation of recombinant HIF-2α. Incubation with recombinant LOX (rLOX) induces HIF-2α carbonylation, which is suppressed by LOX inhibitor treatment, LXG6403 (30µM for 2 hours). **(G)** Kaplan-Meier disease-free survival analysis based on HUWE1 expression in ccRCC patients from TCGA (n = 435). **(H)** Immunoblot analysis showing that inhibition of HECT-domain E3 ligases with heclin rescues HIF-2α protein levels following LOX inhibition, implicating HECT-type ligases in LOX-dependent HIF-2α turnover (LXG6403 30µM for 4 hours; Heclin 20µM for 30 minutes). Survival differences were evaluated using the log-rank (Mantel-Cox) test (G). *P*-values are indicated.

**Supplementary Figure 3.**
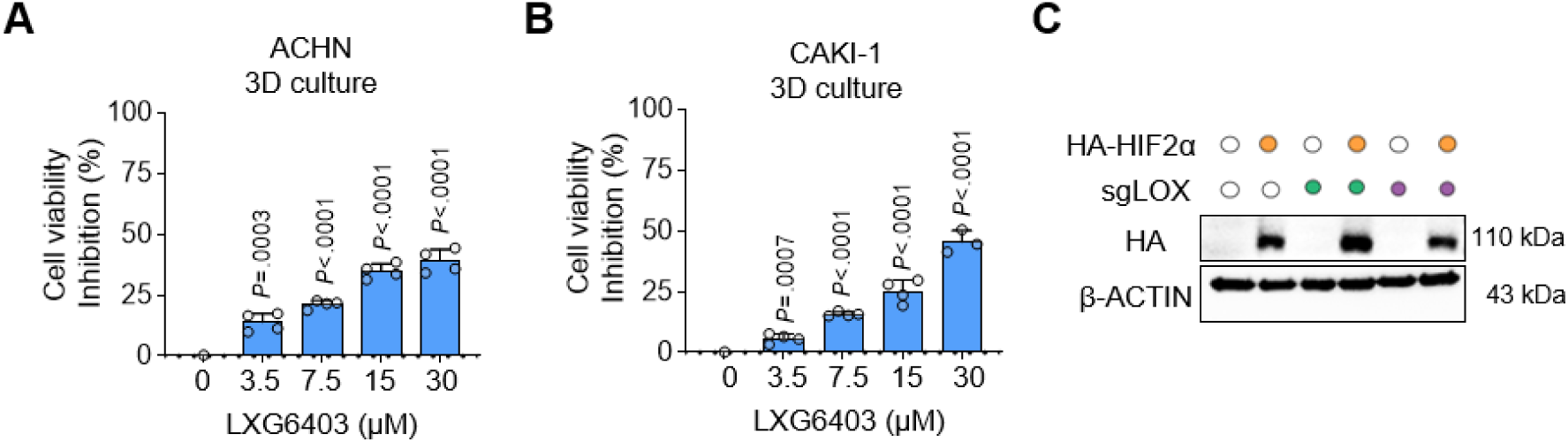
LOX inhibition reduces 3D cell viability in VHL-wildtype ccRCC. (A, **B)** Dose-dependent reduction of 3D viability with LXG6403 treatment in *VHL*-wildtype cells ACHN (A) and CAKI-1 (B) (n = 4 technical replicates). **(C)** Immunoblot validating HA-HIF-2α overexpression in A704 sgLOX cells. Data are shown as mean ± SD. One-way ANOVA with Dunnett’s test was performed to compare more than one group (A, B). *P*-values are indicated.

**Supplementary Figure 4.**
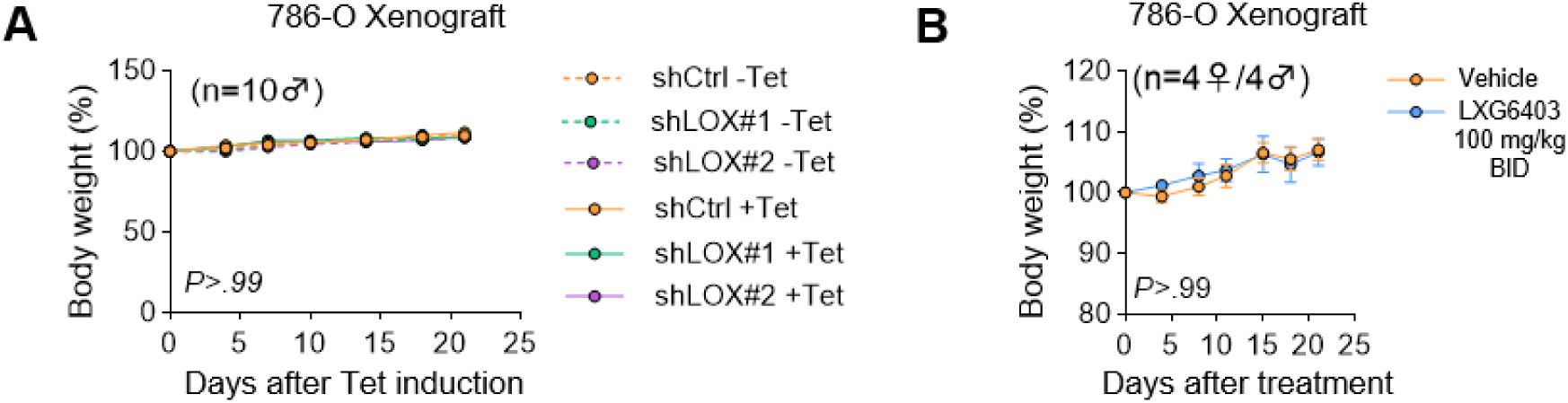
LOX inhibition does not cause toxicity in vivo. (A, **B)** Body weight curves for NSG mice bearing 786-O xenografts with tetracycline-inducible shLOX or control during tetracycline administration (drinking water supplemented with 50 g/L sucrose and 1 g/L tetracycline) (n = 10 mice) (A), and in a sex-balanced 786-O xenograft cohort (n = 4 females, 4 males) treated with vehicle or LXG6403 (100 mg/kg, BID, p.o.) for 21 days (n = 8 NSG mice per group) (B). Data are shown as mean ± SEM. Statistical significance was determined by multiple comparisons with Holm’s correction (A, B). *P*-values are indicated.

**Supplementary Figure 5.**
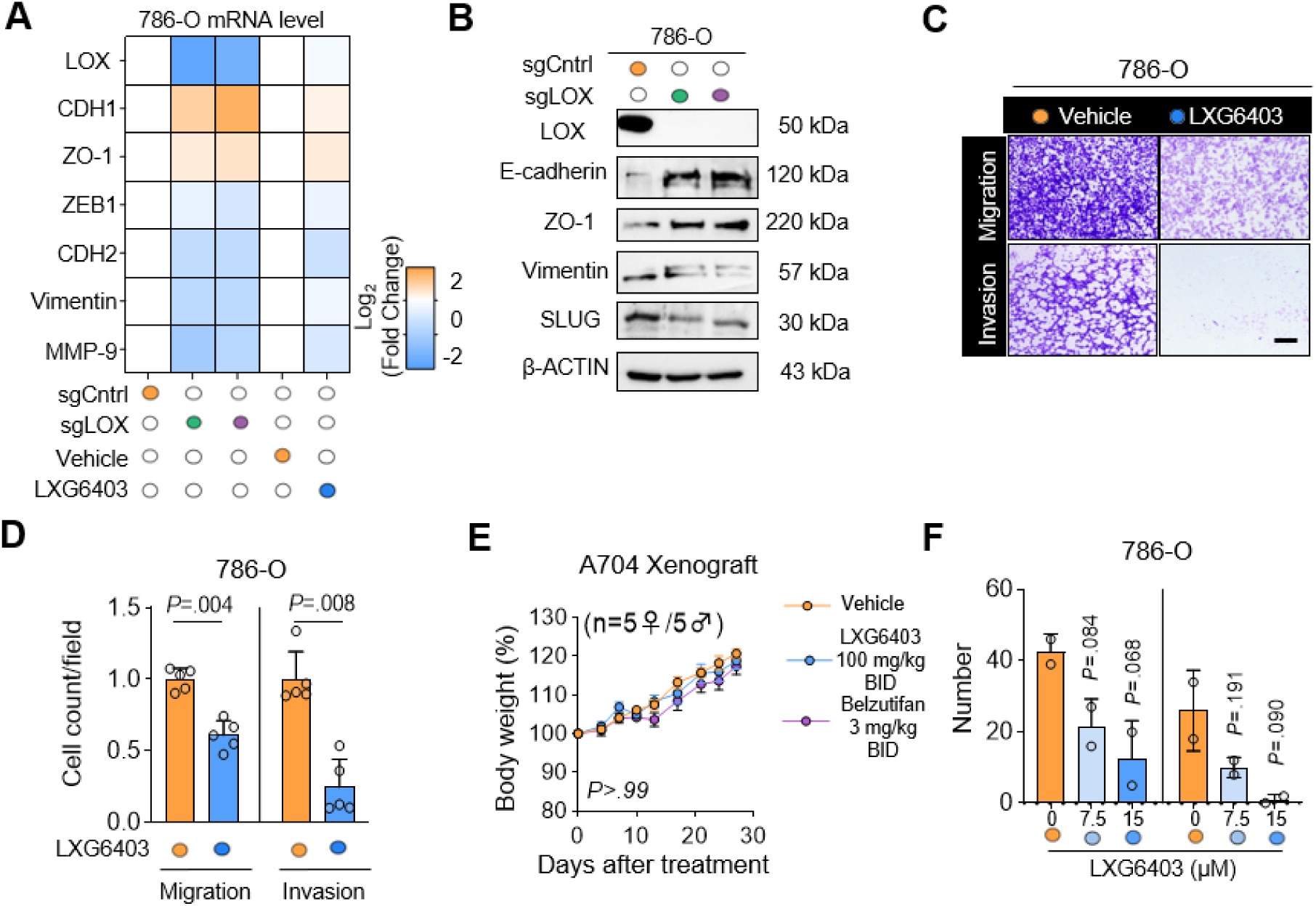
Genetic or pharmacologic inhibition of LOX induces MET-like transition in VHL-mutant ccRCC. **(A)** Heatmap of qRT-PCR analysis of EMT marker changes upon siRNA-mediated LOX knockdown in 786-O cells (n = 2). **(B)** Immunoblot confirming EMT marker changes in sgLOX-expressing 786-O cells. **(C, D)** Transwell migration and invasion assays showing impaired motility in 786-O cells after LOX inhibition (C) and quantification (D) (n = 5 different images). **(E**) Body weight change of mice, showing no significant treatment-related toxicity (n = 5 female/5male NSG mice). **(F)** Quantification of spheres/sub-spheres number in 786-O cells treated with LXG6403 (7.5 and 15 µM) for 21 days (n = 2). Data are shown as mean ± SD from biological experiments. In vivo data are shown as mean ± SEM. Statistical significance was determined by unpaired two-sided Student’s t-test between the two groups (D, F) and multiple comparisons with Tukey’s and Holm’s correction for in vivo data (E). *P*-values are indicated.

**Supplementary Figure 6.**
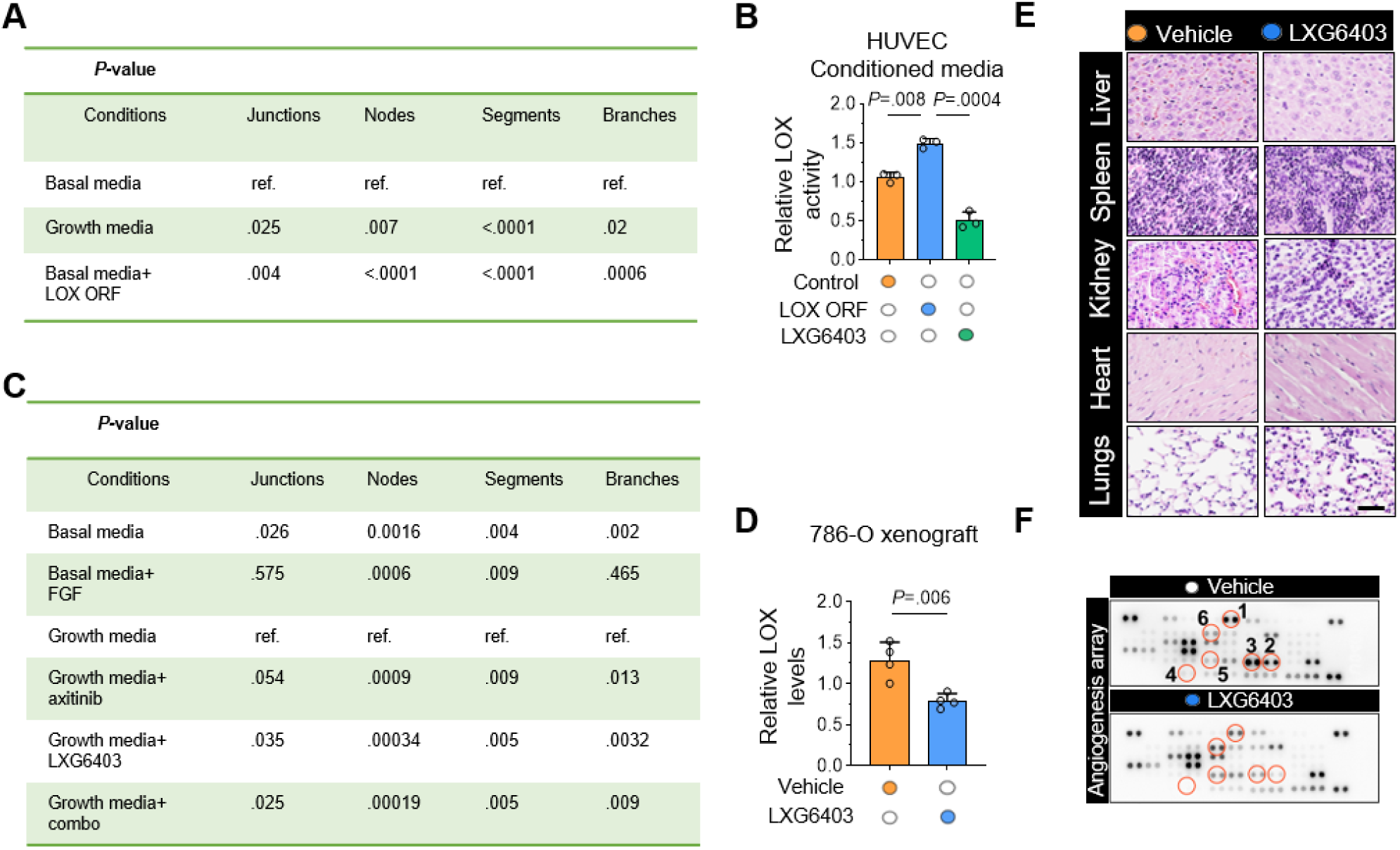
LOX regulates angiogenic activity and hypoxic signaling in ccRCC models. **(A)** Statistical summary reporting *P*-values for HUVEC tubule formation metrics (junctions, nodes, segments, branches) across indicated conditions (Basal media, Growth media, and Basal media+ LOX-conditioned media (CM)). *P*-values derived from one-way ANOVA with Dunnett’s (vs control) and Tukey’s (all pairwise) post hoc tests. **(B)** Relative LOX activity in HUVEC conditioned media (n = 3 technical replicates) **(C)** Statistical summary reporting *P*-values for HUVEC tubule formation metrics (junctions, nodes, segments, branches) across indicated conditions (Basal media, Basal media with Fibroblast growth factor (FGF), Growth media with axitinib, Growth media with LXG6403, and Growth media with combination). *P*-values derived from one-way ANOVA with Dunnett’s (vs control) and Tukey’s (all pairwise) post hoc tests. **(D)** Quantification of serum LOX levels in NSG mice treated with vehicle and LXG6403 (100 mg/kg, p.o, 8h) (n = 4 mice). **(E)** H&E staining of non-tumor organs showed no detectable histopathological abnormalities following LXG6403 treatment (n = 4 mice). **(F)** Angiogenic protein array blot (1: Angiopoietin-1, 2: PDGF-AA, 3: PDGF-AB/-BB, 4: VEGF, 5: Endostatin, 6: Platelet factor 4), (n = 2 mice). Data are shown as mean ± SD. Statistical significance was determined by unpaired two-sided Student’s t-test between the two groups (B, D). *P*-values are indicated.

**Supplementary Figure 7.**
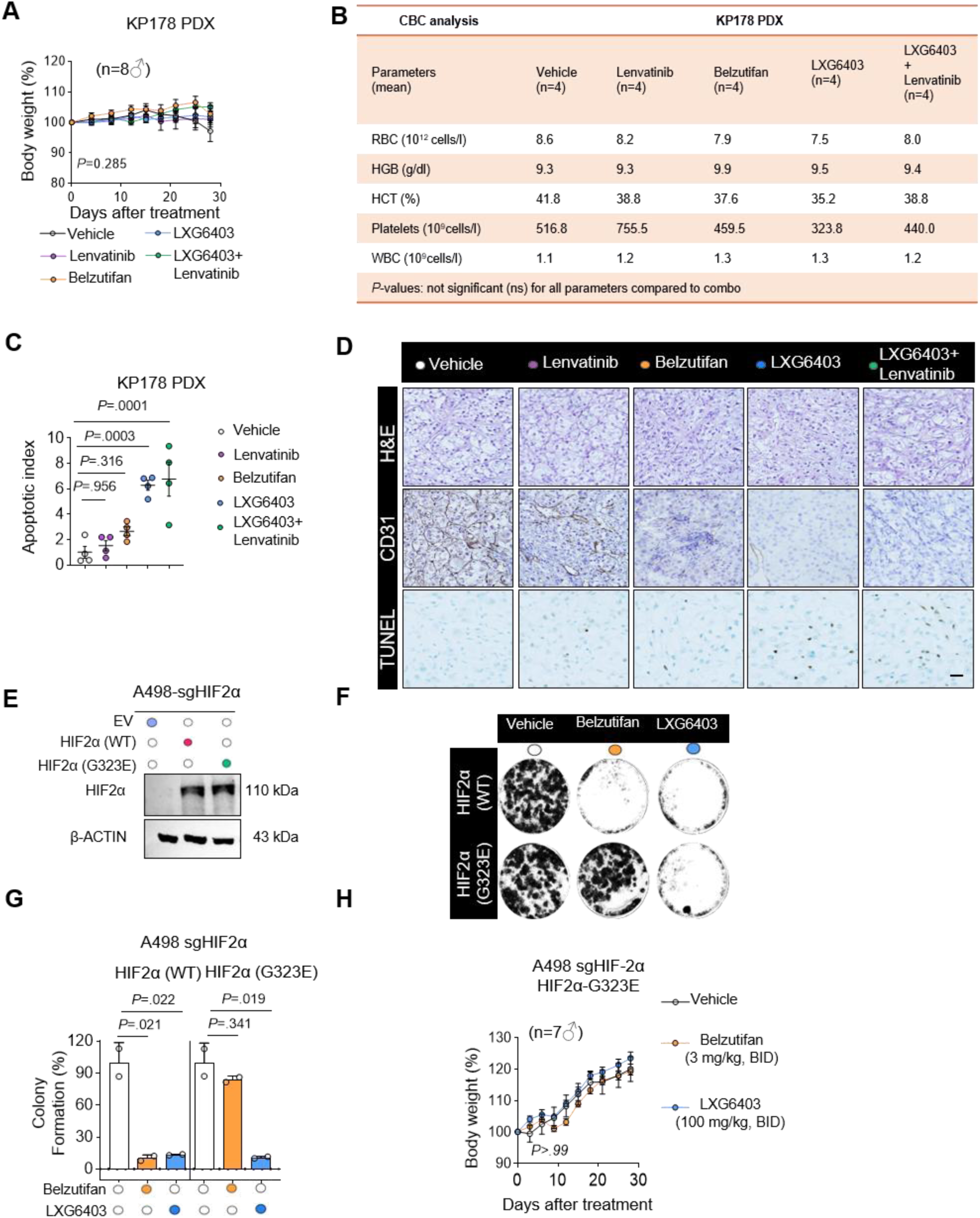
LOX inhibition in combination with lenvatinib does not cause toxicity while inducing apoptosis, and the validation of HIF-2α expression upon wildtype or G323E mutant HIF-2α overexpression. **(A)** Percent body weight change in the KP78 PDX study: vehicle, lenvatinib (10 mg/kg, QD, p.o), LXG6403 (75 mg/kg, BID, p.o.), and combination for 28 days (n = 8 male NSG mice) and immunoblot from KP178 ccRCC PDX tumor lysate showing high levels of LOX and VEGFA, β-actin as loading control. **(B)** Complete blood counts (RBC, HGB, HCT, WBC, PLT) from KP178 PDX study demonstrating no hematologic toxicity (n = 4 mice). **(C)** Quantification of apoptotic index from TUNEL-stained tumor sections from (A) (n = 4 tumors). **(D)** Representative images of H&E, CD31 and TUNEL staining in tumor sections from (A) (n = 4 tumors). **(E)** Immunoblot showing overexpression of HIF-2α wildtype (WT) and HIF-2α G323E in sgHIF-2α-expressing A498 cells. **(F)** Colony formation assay showing reduced clonogenic potential upon LOX (15 μM) or belzutifan (1μM) treatment in sgHIF-2α-expressing A498 cells overexpressed either with HIF-2α WT or HIF-2α G323E (n = 2). **(G)** Quantification of (F). **(H)** Percent body weight change in A498 sgHIF-2α HIF-2α-G323E xenograft cohort treated with vehicle, belzutifan (3 mg/kg, BID, p.o.), or LXG6403 (100 mg/kg, BID, p.o.) for 28 days (n = 7 male mice). Data are shown as mean ± SD from biological experiments. In vivo data are shown as mean ± SEM. Statistical significance was determined by unpaired two-sided Student’s T-test (B) or two-way ANOVA with Dunett’s (C, G), and Holm’s (A, H) corrections. *P*-values are indicated.

---

**Supplementary Table 1.** Clinicopathologic characteristics of the ccRCC tissue microarrays (TMAs)

| **Entire TMA Cohort, n** |  | **204** |
| --- | --- | --- |
| **Histologic Grade, n (%)** |  |  |
| *I* |  | 16 (7.8%) |
| *II* |  | 90 (44.1%) |
| *III* |  | 83 (40.7%) |
| *IV* |  | 15 (7.4%) |
| *Clinical subset\*, n* |  | 65 out of 204 |
| **Age, Years, Median (Range)** |  | 60 (23-87) |
| **Sex, n (%)** |  |  |
| *Male* |  | 44 (67.7%) |
| *Female* |  | 21 (32.3%) |
| **Pathologic T Stage, n (%)** |  |  |
| *pT1* |  | 38 (58.5%) |
| *pT2* |  | 0 (0%) |
| *pT3* |  | 26 (40%) |
| *pT4* |  | 1 (1.5%) |
| *Clinical annotations (age, sex, and pathologic t stage) were available for 65 of the 204 patients included in the TMA cohort. The remaining cases were reported previously<sup>61</sup>. |  |  |

**Supplementary Table 2.** List of antibodies used in WB, IP, and IF.

| *List of antibodies used in Western blot (WB), Proximity ligation assay (PLA), immunohistochemistry (IHC), immunoprecipitation (IP) and chromatin immunoprecipitation (ChIP) experiments.* |  |  |  |  |  |
| --- | --- | --- | --- | --- | --- |
| Antibody | Provider | Catalog number | WB dilution | IP dilution | PLA/IHC/ChIP dilution |
| LOX | Sigma-Aldrich | L4794<br>(RRID:AB_1840993) | 1:1000 | 1:250 | 1:100 (PLA),<br>1:600 (IHC) |
| HIF-2α | Novus | NB100-122<br>(RRID:AB_10002593) | 1:1000 | 1:100 | 1:600 (IHC),<br>1:100 (ChIP) |
| HIF-2α | Santa Cruz | sc-46691<br>(RRID:AB_627523) | | | 1:100 (PLA) |
| HA | Invitrogen | 26183<br>(RRID:AB_2533049) | 1:7500 |  | - |
| FLAG | Proteintech | 80010-1-RR<br>(RRID:AB_2882940) | 1:7500 | 1:250 | - |
| β-ACTIN | Sigma | A2228<br>(RRID:AB_476697) | 1:10000 | - | - |
| UBIQUITIN | Cell Signaling | 58395<br>(RRID:AB_3075532) | 1:2000 | - | - |
| STREPTAVIDIN-HRP | Thermo Fischer | EMDA5441<br>(RRID:AB_2619743) | 1:7500 | - | - |
| VHL | Cell signaling | 68547<br>(RRID:AB_271627) | 1:1000 | - | - |
| E-CADHERIN | BD Biosciences | BD610181<br>(RRID:AB_397580) | 1:1000 | - | - |
| ZO-1 | Cell signaling | 5406<br>(RRID:AB_1904187) | 1:1000 | - | - |
| VIMENTIN | Cell signaling | 5741<br>(RRID:AB_10695459) | 1:1000 | - | - |
| ZEB-1 | Santa Cruz | sc-515797<br>(RRID:AB_2934316) | 1:1000 | - | - |
| SLUG | Abcam | ab27568<br>(RRID:AB_777968) | 1:1000 | - | - |
| GAPDH | Cell signaling | 2118<br>(RRID:AB_561053) | 1:1000 | - | - |
| α-TUBULIN | Santa Cruz | sc-32293<br>(RRID:AB_628412) | 1:1000 | - | - |
| LAMIN B1 | Santa Cruz | sc-374015<br>(RRID:AB_10947408) | 1:1000 | - | - |
| PDGFβ | Cell Signaling | 3169<br>(RRID:AB_2162497) | 1:1000 | - | - |
| p-PDGFβ | Cell Signaling | 3166<br>(RRID:AB_331365) | 1:1000 | - | - |
| FGFR1 | Cell Signaling | 9740<br>(RRID:AB_11178519) | 1:1000 | - | - |
| p-FGFR1 | Cell Signaling | 52928<br>(RRID:AB_2799424) | 1:1000 | - | - |
| VEGFR2 | Cell Signaling | 2472<br>(RRID:AB_33111) | 1:1000 | - | - |
| p-VEGFR2 | Cell Signaling | 2478<br>(RRID:AB_331377) | 1:1000 | - | - |
| ANG-1 | Proteintech | 27093-1-AP<br>(RRID:AB_3085925) | 1:1000 | - | - |
| VEGFA | Cell Signaling | 50661(RRID:<br>AB_3751965) | 1:1000 | - | - |
| SRC | Cell Signaling | 2109<br>(RRID:AB_2106059) | 1:1000 | - | - |
| p-SRC | Cell Signaling | 2101<br>(RRID:AB_331697) | 1:1000 | - | - |
| FAK | Cell Signaling | 3285<br>(RRID:AB_2269034) | 1:1000 | - | - |
| AKT | Santa Cruz | 9272<br>(RRID:AB_329827) | 1:1000 | - | - |
| p-AKT | Cell Signaling | 13038<br>(RRID:AB_2629447) | 1:1000 | - | - |
| HRP-LINKED ANTI-MOUSE IGG | Cell Signaling | 7076<br>(RRID:AB_330924) | 1:5000 | - | - |
| HRP-LINKED ANTI-RABBIT IGG | Cell Signaling | 7074<br>(RRID:AB_2099233) | 1:5000 | - | - |

**Supplementary Table 3.**
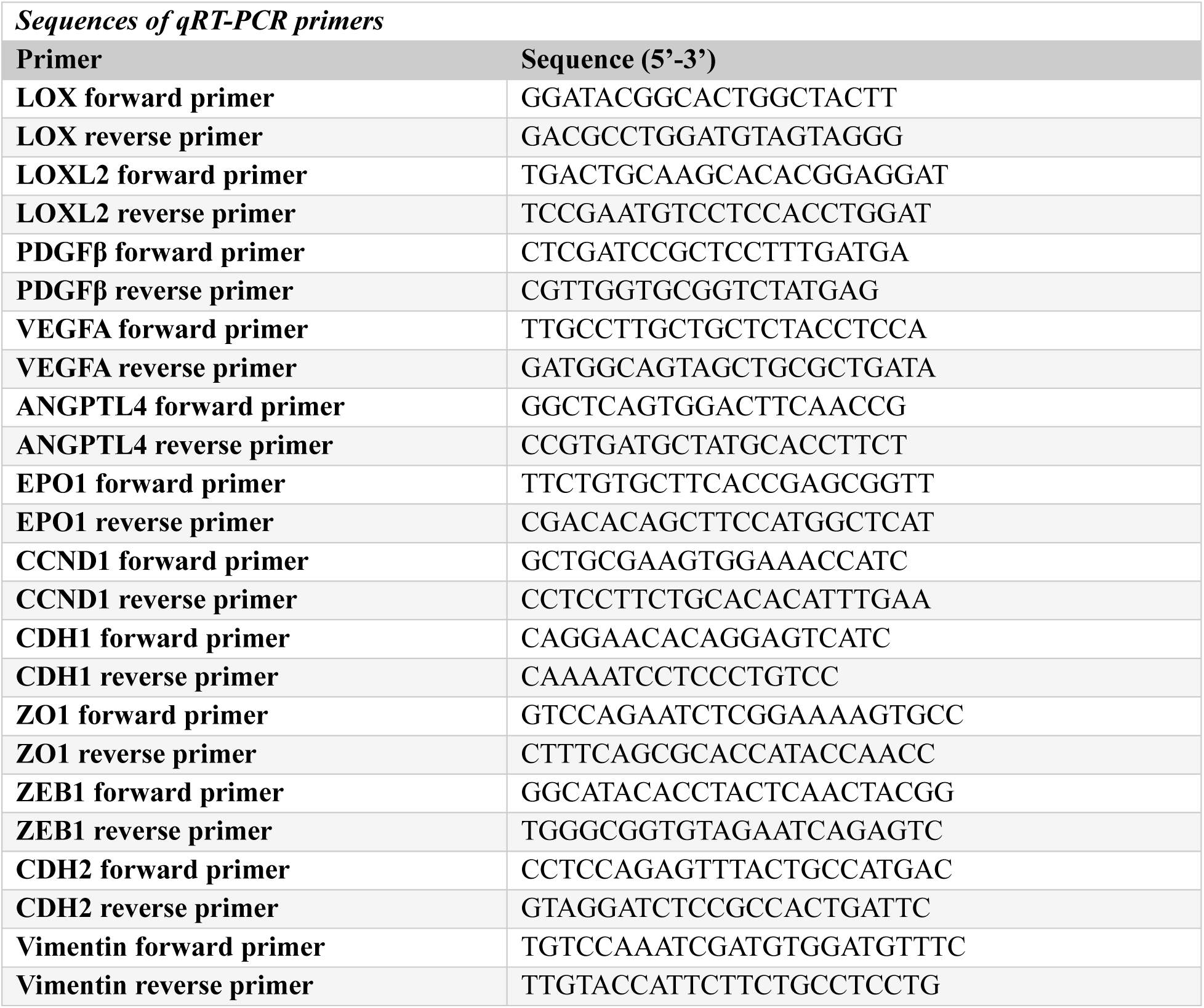
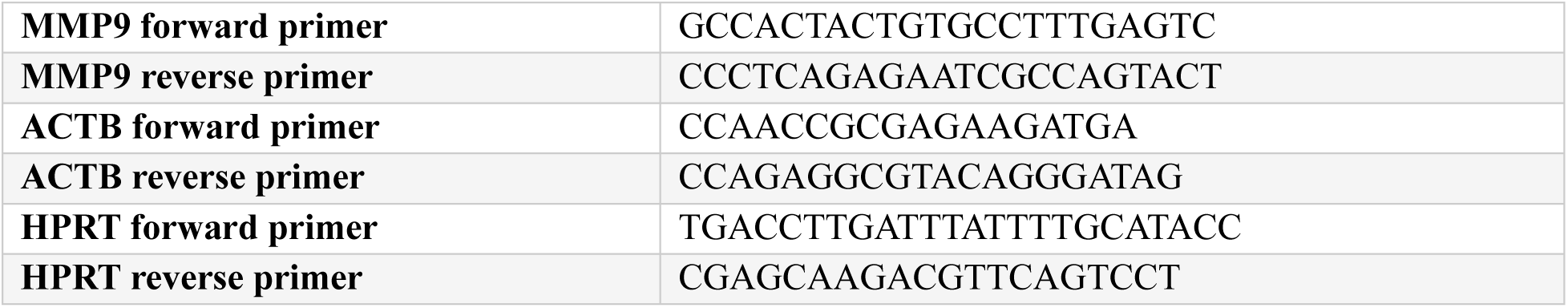
Primer sequences used in qRT-PCR.

**Supplementary Table 4.**
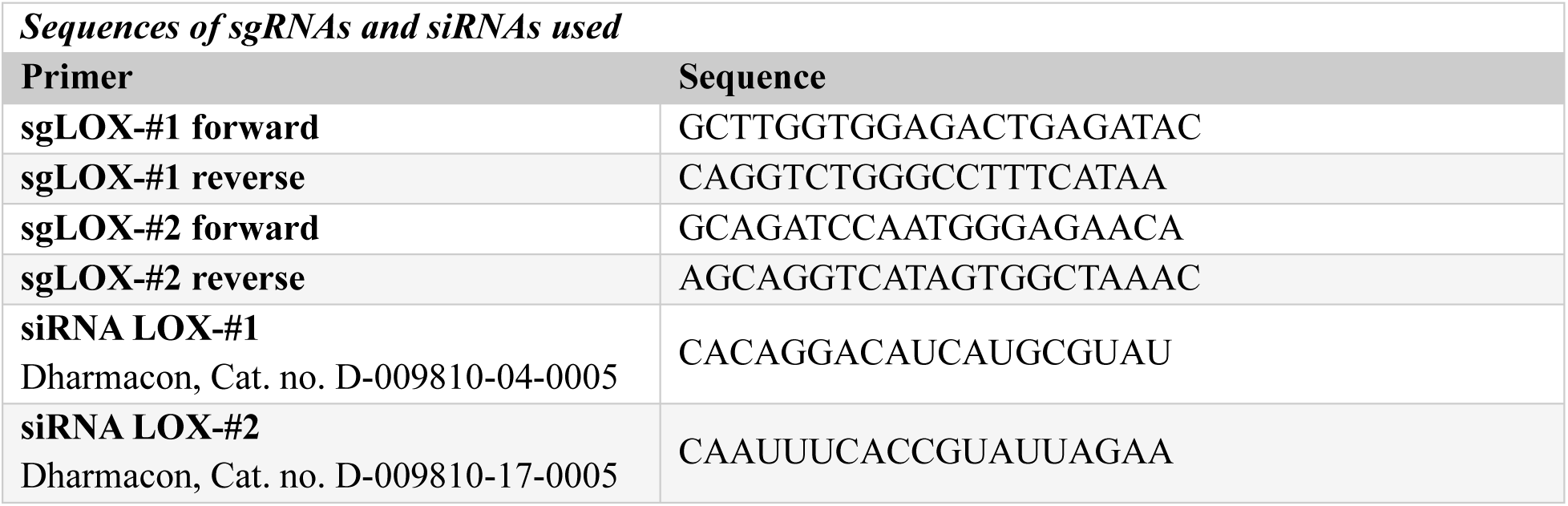
siRNA and sgRNA sequences.

**Supplementary Table 5.**
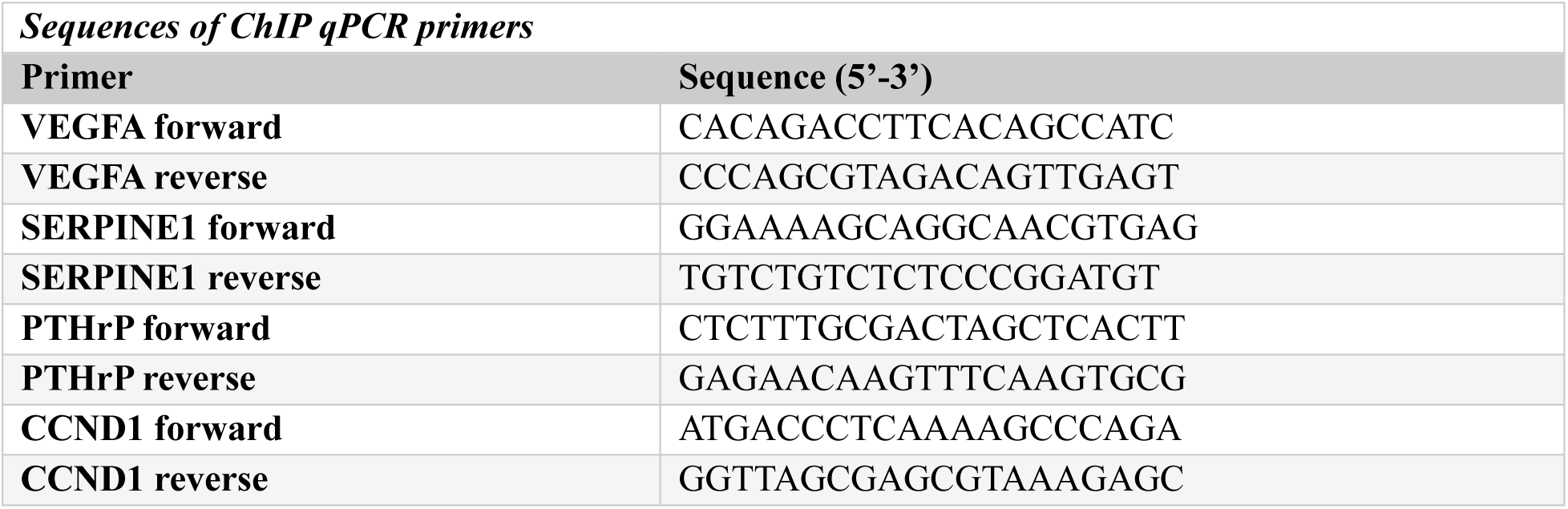
ChIP-qPCR primers.

## References

1. National Cancer Institute Surveillance, E. a. E. R. P. Cancer stat facts: kidney and renal pelvis cancer https://seer.cancer.gov/statfacts/html/kidrp.html (2025).

2. Sung, H., et al. Global cancer statistics 2020: GLOBOCAN estimates of incidence and mortality worldwide for 36 cancers in 185 countries. CA: a cancer journal for clinicians 71, 209–249 (2021).

3 Moch, H., Cubilla, A. L., Humphrey, P. A., Reuter, V. E. & Ulbright, T. M. The 2016 WHO classification of tumours of the urinary system and male genital organs part A: renal, penile, and testicular tumours. European urology 70, 93–105 (2016).

4 Powles, T. et al. Renal cell carcinoma: ESMO Clinical Practice Guideline for diagnosis, treatment and follow-up. Ann Oncol 35, 692–706 (2024). 10.1016/j.annonc.2024.05.537

5 Motzer, R. J. et al. NCCN Guidelines® Insights: Kidney Cancer, Version 2.2024: Featured Updates to the NCCN Guidelines. Journal of the National Comprehensive Cancer Network 22, 4–16 (2024). 10.6004/jnccn.2024.0008

6 Brugarolas, J. Molecular Genetics of Clear-Cell Renal Cell Carcinoma. Journal of Clinical Oncology 32, 1968–1976 (2014). 10.1200/jco.2012.45.2003

7 Riazalhosseini, Y. & Lathrop, M. Precision medicine from the renal cancer genome. Nat Rev Nephrol 12, 655–666 (2016). 10.1038/nrneph.2016.133

8 Latif, F. et al. Identification of the von Hippel-Lindau disease tumor suppressor gene. Science 260, 1317–1320 (1993).

9 Madrigal, A. et al. Single-cell gene programs define subtype identity and metastatic trajectories in renal cell carcinoma. medRxiv, 2026.2007.2014.26357682 (2026). 10.64898/2026.07.14.26357682

10 Gerlinger, M. et al. Intratumor heterogeneity and branched evolution revealed by multiregion sequencing. N Engl J Med 366, 883–892 (2012). 10.1056/NEJMoa1113205

11 Ahluwalia, P. et al. Prognostic and therapeutic implications of extracellular matrix associated gene signature in renal clear cell carcinoma. Scientific Reports 11, 75–61 (2021). 10.1038/s41598-021-86888-7

12 Madrigal, A. et al. Abstract PR005: An atlas of cellular heterogeneity in primary and metastatic renal cell carcinomas. Cancer Research 83, PR005–PR005 (2023). 10.1158/1538-7445.Kidney23-pr005

13 Miheecheva, N. et al. Multiregional single-cell proteogenomic analysis of ccRCC reveals cytokine drivers of intratumor spatial heterogeneity. Cell Rep 40, 111–180 (2022). 10.1016/j.celrep.2022.111180

14 Mandriota, S. J. et al. HIF activation identifies early lesions in VHL kidneys: Evidence for site-specific tumor suppressor function in the nephron. Cancer Cell 1, 459–468 (2002). 10.1016/S1535-6108(02)00071-5

15 Choueiri, T. K. et al. Nivolumab plus Cabozantinib versus Sunitinib for Advanced Renal-Cell Carcinoma. New England Journal of Medicine 384, 829–841 (2021). doi:10.1056/NEJMoa2026982

16 Thakur, A. et al. Recent advances and future directions on small molecule VEGFR inhibitors in oncological conditions. European Journal of Medicinal Chemistry 272, 116472 (2024). 10.1016/j.ejmech.2024.116472

17 Sharma, R. et al. Determinants of resistance to VEGF-TKI and immune checkpoint inhibitors in metastatic renal cell carcinoma. Journal of Experimental & Clinical Cancer Research 40, 186 (2021).

18 Chen, W. et al. Targeting renal cell carcinoma with a HIF-2 antagonist. Nature 539, 112–117 (2016). 10.1038/nature19796

19 Lawson, K. V. et al. Abstract A012: Discovery and characterization of Casdatifan (AB521), a clinical-stage, potent, and selective Hypoxia-Inducible Factor (HIF)-2α inhibitor. Molecular Cancer Therapeutics 23, A012 (2024). 10.1158/1538-8514.Cancerchem24-a012

20 Lu, J. et al. Abstract 6330: NKT2152: A highly potent HIF2α inhibitor and its therapeutic potential in solid tumors beyond ccRCC. Cancer Research 82, 6330–6330 (2022). 10.1158/1538-7445.Am2022-6330

21 Xu, R. et al. 3-[(1S,2S,3R)-2,3-Difluoro-1-hydroxy-7-methylsulfonylindan-4-yl]oxy-5-fluorobenzonitrile (PT2977), a Hypoxia-Inducible Factor 2α (HIF-2α) Inhibitor for the Treatment of Clear Cell Renal Cell Carcinoma. Journal of Medicinal Chemistry 62, 6876–6893 (2019). 10.1021/acs.jmedchem.9b00719

22 Schweickert, P. G. et al. Casdatifan (AB521) is a novel and potent allosteric small molecule inhibitor of protumourigenic HIF-2α dependent transcription. Br J Pharmacol 182, 4147–4167 (2025). 10.1111/bph.70075

23 Deeks, E. D. Belzutifan: first approval. Drugs 81, 1921–1927 (2021).

24 Jonasch, E. et al. 1690O NKT2152, a novel oral HIF-2 inhibitor, in participants (pts) with previously treated advanced clear cell renal carcinoma (accRCC): Preliminary results of a phase I/II study. Annals of Oncology 35, S1011–S1012 (2024). 10.1016/j.annonc.2024.08.1783

25 Natarajan, V., Satalkar, V., Gumbart, J. C. & Torres, M. Molecular Dynamics Reveals Altered Interactions between Belzutifan and HIF-2 with Natural Variant G323E or Proximal Phosphorylation at T324. ACS Omega 9, 37843–37855 (2024). 10.1021/acsomega.4c03777

26 Courtney, K. D. et al. HIF-2 complex dissociation, target inhibition, and acquired resistance with PT2385, a first-in-class HIF-2 inhibitor, in patients with clear cell renal cell carcinoma. Clinical Cancer Research 26, 793–803 (2020).

27 Stransky, L. A. et al. Sensitivity of VHL mutant kidney cancers to HIF2 inhibitors does not require an intact p53 pathway. Proceedings of the National Academy of Sciences 119, e2120403119 (2022).

28 James B. Brugarolas, H. H., Tao Wang. Methods of identifying and treating patients with HIF-2 inhibitor resistance. USA patent (2023).

29 Kagan, H. M. & Li, W. Lysyl oxidase: properties, specificity, and biological roles inside and outside of the cell. J Cell Biochem 88, 660–672 (2003). 10.1002/jcb.10413

30 Pinnell, S. R. & Martin, G. R. The cross-linking of collagen and elastin: enzymatic conversion of lysine in peptide linkage to alpha-aminoadipic-delta-semialdehyde (allysine) by an extract from bone. Proceedings of the National Academy of Sciences 61, 708–716 (1968). doi:10.1073/pnas.61.2.708

31 Madrigal A., K. M., Mehrjoo Z., Nishimura T., Saatci O., Osakwe A., Moslemi E., Glennon K. I., Dankner M., Maritan S., Kuasne H., Pilon V., Monast A., Soytas M., Arseneault M., Oikonomopoulos S., Harutyunyan A., Lu T., Rayes R., Soto L.M., Hernandez-Corchado A., Spicer J.D., Petrecca K., Siegel P., Park M., Ragoussis J., Sahin O., Brimo F., Tanguay S., Riazalhosseini Y., Najafabadi H.S. Single-cell gene programs define subtype identity and metastatic trajectories in renal cell carcinoma *Preprint at bioRxiv* (2026).

32 Guo, L., An, T., Wan, Z., Huang, Z. & Chong, T. SERPINE1 and its co-expressed genes are associated with the progression of clear cell renal cell carcinoma. BMC Urol 23, 43 (2023). 10.1186/s12894-023-01217-6

33 McEvoy, C. M. et al. Single-cell profiling of healthy human kidney reveals features of sex-based transcriptional programs and tissue-specific immunity. Nat Commun 13, 7634 (2022). 10.1038/s41467-022-35297-z

34 Su, C. et al. Single-Cell RNA Sequencing in Multiple Pathologic Types of Renal Cell Carcinoma Revealed Novel Potential Tumor-Specific Markers. Front Oncol 11, 719564 (2021). 10.3389/fonc.2021.719564

35 Wang, V., Davis, D. A. & Yarchoan, R. Identification of functional hypoxia inducible factor response elements in the human lysyl oxidase gene promoter. Biochemical and Biophysical Research Communications 490, 480–485 (2017). 10.1016/j.bbrc.2017.06.066

36 Pez, F. et al. The HIF-1-inducible lysyl oxidase activates HIF-1 via the Akt pathway in a positive regulation loop and synergizes with HIF-1 in promoting tumor cell growth. Cancer Res 71, 1647–1657 (2011). 10.1158/0008-5472.Can-10-1516

37 Lucero, H. A. et al. Lysyl oxidase oxidizes cell membrane proteins and enhances the chemotactic response of vascular smooth muscle cells. J Biol Chem 283, 24103–24117 (2008). 10.1074/jbc.M709897200

38 Kamadurai, H. B. et al. Insights into ubiquitin transfer cascades from a structure of a UbcH5B approximately ubiquitin-HECT(NEDD4L) complex. Mol Cell 36, 1095–1102 (2009). 10.1016/j.molcel.2009.11.010

39 Mund, T., Lewis, M. J., Maslen, S. & Pelham, H. R. Peptide and small molecule inhibitors of HECT-type ubiquitin ligases. Proc Natl Acad Sci U S A 111, 16736–16741 (2014). 10.1073/pnas.1412152111

40 Levental, K. R. et al. Matrix Crosslinking Forces Tumor Progression by Enhancing Integrin Signaling. Cell 139, 891–906 (2009). 10.1016/j.cell.2009.10.027

41 Saatci, O. et al. Targeting lysyl oxidase (LOX) overcomes chemotherapy resistance in triple negative breast cancer. Nat Commun 11, 2416 (2020). 10.1038/s41467-020-16199-4

42 Cetin, M. et al. A highly potent bi-thiazole inhibitor of LOX rewires collagen architecture and enhances chemoresponse in triple-negative breast cancer. Cell Chem Biol 31, 1926–1941.e1911 (2024). 10.1016/j.chembiol.2024.06.012

43 Yan, Q., Bartz, S., Mao, M., Li, L. & Kaelin, W. G., Jr. The hypoxia-inducible factor 2alpha N-terminal and C-terminal transactivation domains cooperate to promote renal tumorigenesis in vivo. Mol Cell Biol 27, 2092–2102 (2007). 10.1128/mcb.01514-06

44 Mani, S. A. et al. The epithelial-mesenchymal transition generates cells with properties of stem cells. Cell 133, 704–715 (2008). 10.1016/j.cell.2008.03.027

45 Ayers, G. D. et al. Volume of preclinical xenograft tumors is more accurately assessed by ultrasound imaging than manual caliper measurements. J Ultrasound Med 29, 891–901 (2010). 10.7863/jum.2010.29.6.891

46 Chappell, J. C., Payne, L. B. & Rathmell, W. K. Hypoxia, angiogenesis, and metabolism in the hereditary kidney cancers. J Clin Invest 129, 442–451 (2019). 10.1172/JCI120855

47 Choueiri, T. K. & Kaelin, W. G., Jr. Targeting the HIF2-VEGF axis in renal cell carcinoma. Nat Med 26, 1519–1530 (2020). 10.1038/s41591-020-1093-z

48 Gerhardt, H. et al. VEGF guides angiogenic sprouting utilizing endothelial tip cell filopodia. J Cell Biol 161, 1163–1177 (2003). 10.1083/jcb.200302047

49 Cotta, B. H. et al. Current Landscape of Genomic Biomarkers in Clear Cell Renal Cell Carcinoma. Eur Urol 84, 166–175 (2023). 10.1016/j.eururo.2023.04.003

50 Scelo, G. et al. Variation in genomic landscape of clear cell renal cell carcinoma across Europe. Nature Communications 5, 5135 (2014). 10.1038/ncomms6135

51 van der Mijn, J. C. et al. The genomic landscape of metastatic clear cell renal cell carcinoma after systemic therapy. Mol Oncol 16, 2384–2395 (2022). 10.1002/1878-0261.13204

52 Chen, W. J. et al. Single-cell RNA-seq integrated with multi-omics reveals SERPINE2 as a target for metastasis in advanced renal cell carcinoma. Cell Death Dis 14, 30 (2023). 10.1038/s41419-023-05566-w

53 Kaelin Jr, W. G. The von Hippel–Lindau tumour suppressor protein: O2 sensing and cancer. Nature Reviews Cancer 8, 865–873 (2008).

54 Gossage, L., Eisen, T. & Maher, E. R. VHL, the story of a tumour suppressor gene. Nat Rev Cancer 15, 55–64 (2015). 10.1038/nrc3844

55 Linehan, W. M. et al. The Metabolic Basis of Kidney Cancer. Cancer Discovery 9, 1006–1021 (2019). 10.1158/2159-8290.Cd-18-1354

56 Linehan, W. M., Rubin, J. S. & Bottaro, D. P. VHL loss of function and its impact on oncogenic signaling networks in clear cell renal cell carcinoma. Int J Biochem Cell Biol 41, 753–756 (2009). 10.1016/j.biocel.2008.09.024

57 Patard, J. J. et al. Absence of VHL gene alteration and high VEGF expression are associated with tumour aggressiveness and poor survival of renal-cell carcinoma. Br J Cancer 101, 1417–1424 (2009). 10.1038/sj.bjc.6605298

58 Choueiri, T. K. et al. The role of aberrant VHL/HIF pathway elements in predicting clinical outcome to pazopanib therapy in patients with metastatic clear-cell renal cell carcinoma. Clin Cancer Res 19, 5218–5226 (2013). 10.1158/1078-0432.Ccr-13-0491

59 Cho, H. & Kaelin, W. G. in Cold Spring Harbor symposia on quantitative biology. 113–121 (Cold Spring Harbor Laboratory Press).

60 Wiesener, M. S. et al. Widespread, hypoxia-inducible expression of HIF-2α in distinct cell populations of different organs. The FASEB Journal 17, 271–273 (2003).

61 Xu, B. et al. Enhancer of Zeste Homolog 2 Expression Is Associated With Metastasis and Adverse Clinical Outcome in Clear Cell Renal Cell Carcinoma: A Comparative Study and Review of the Literature. Archives of Pathology & Laboratory Medicine 137, 1326–1336 (2013). 10.5858/arpa.2012-0525-OA

62 Liberzon, A. et al. The molecular signatures database hallmark gene set collection. Cell systems 1, 417–425 (2015).

63 Lun, A. T., McCarthy, D. J. & Marioni, J. C. A step-by-step workflow for low-level analysis of single-cell RNA-seq data with Bioconductor. F1000Research 5, 2122 (2016).

## SUPPLEMENTARY REFERENCES

1. Mishra, R.R., et al. Reactivation of cAMP Pathway by PDE4D Inhibition Represents a Novel Druggable Axis for Overcoming Tamoxifen Resistance in ER-positive Breast Cancer. Clin Cancer Res 24, 1987–2001 (2018).

2. Assidicky, R., et al. Targeting HIF1-alpha/miR-326/ITGA5 axis potentiates chemotherapy response in triple-negative breast cancer. Breast Cancer Res Treat 193, 331–348 (2022).

3. Carpentier, G., et al. Angiogenesis Analyzer for ImageJ - A comparative morphometric analysis of “Endothelial Tube Formation Assay” and “Fibrin Bead Assay”. Sci Rep 10, 11568 (2020).

4. Saatci, O., et al. Targeting lysyl oxidase (LOX) overcomes chemotherapy resistance in triple negative breast cancer. Nat Commun 11, 2416 (2020).

